# Single-cell proteomics reveals downregulation of TMSB4X to drive actin release for stereocilia assembly

**DOI:** 10.1101/727412

**Authors:** Ying Zhu, Mirko Scheibinger, Daniel C. Ellwanger, Jocelyn F. Krey, Dongseok Choi, Ryan T. Kelly, Stefan Heller, Peter G. Barr-Gillespie

## Abstract

Hearing and balance rely on small sensory hair cells that reside in the inner ear. To explore dynamic changes in the abundant proteins present in differentiating hair cells, we used nanoliter-scale shotgun mass spectrometry of single cells, each ∼1 picoliter, from utricles of embryonic day 15 chickens. We identified unique constellations of proteins or protein groups from presumptive hair cells and from progenitor cells. The single-cell proteomes enabled the *de novo* reconstruction of a developmental trajectory. Inference of protein expression dynamics revealed that the actin monomer binding protein thymosin β4 (TMSB4X) was present in progenitors but dropped precipitously during hair-cell differentiation. Complementary single-cell transcriptome profiling showed downregulation of *TMSB4X* mRNA during maturation of hair cells. We propose that most actin is sequestered by TMSB4X in progenitor cells, but upon differentiation to hair cells, actin is released to build the sensory hair bundle.

## Introduction

Hair cells, the sensory cells of the inner ear, carry out a finely orchestrated construction of an elaborate actin cytoskeleton during differentiation. Progenitors of vestibular hair cells, the supporting cells (Roberson et al., 1992), have an unremarkable actin cytoskeleton. By contrast, differentiating hair cells express a wide array of actin associated proteins, including crosslinkers, membrane-to-actin linkers, and capping molecules, and use them to rapidly assemble mechanically sensitive hair bundles on their apical surfaces (Shin et al., 2013; Ellwanger et al., 2018). Hair bundles consist of ∼100 stereocilia each, filled with filamentous actin (F-actin) and arranged in multiple rows of increasing length; by maturity, stereocilia contain >90% of the F-actin in a hair cell (Tilney and Tilney, 1988).

Along the axis of the chicken cochlea, stereocilia systematically decrease in maximum height (from >5 to 1.5 µm) and increase in number per cell (from 30 to 300) as the frequency encoded increases (Tilney et al., 1992). Despite these changes, a quantitative analysis suggested that within experimental error, hair cells use the same amount of actin to build these disparate hair bundles (Tilney and Tilney, 1988). There is little evidence of substantially increased expression of actin genes during hair cell differentiation (Ellwanger et al., 2018), suggesting that hair cells use the existing monomeric actin that is available during differentiation to build a bundle (Tilney and Tilney, 1988).

The number of hair cells in the chick utricle increases 15-fold from embryonic day 7 (E7) to posthatch day 2, with hair-cell production peaking at E12 (Goodyear et al., 1999). Because of this asynchronous development, at any given age, cells are in distinct states along the pathway from progenitors to hair cells. While bulk sampling of cells averages over these developmental distinctions, sampling and analyzing individual cells of the E15 chick utricle allowed for transcriptomics examination of a large portion of the developmental trajectory for forming hair cells from supporting cells (Ellwanger et al., 2018), and should allow for a corresponding trajectory analysis using proteomics.

We sought to understand how supporting cells, with their modest F-actin cytoskeletons, could transform rapidly into stereocilia-endowed hair cells without significant upregulation of actin gene transcription. We used a highly sensitive single-cell proteomics approach to assess the concentrations of the abundant proteins in cells isolated from the E15 chick utricle. We found that hair cells were readily distinguished from supporting cells based on only 50-75 proteins identified in each. Notably, the actin monomer binding protein thymosin β4 (TMSB4X) was abundant in supporting cells but not in hair cells, and its expression decreased as progenitors developed into hair cells, which we revealed using trajectory analysis based on the proteomics data. Single-cell RNA-seq (scRNA-seq) analysis showed that *TMSB4X* transcripts are downregulated when transcription of *ATOH1*, a key regulator of hair cell differentiation, is activated. Thus, upon differentiation to hair cells, we predict that most of the monomeric actin is made available for hair-bundle assembly.

## Results

### Single-cell proteomics applied to E15 chick utricle hair cells

Recent reports demonstrated that extensive shotgun mass spectrometry characterization of the proteins of single cells is possible when samples are processed in nanoliter-scale volumes (Zhu et al., 2018b; Zhu et al., 2018c; Zhu et al., 2018a; Zhu et al., 2018d). About 700 proteins or protein groups could be detected from a single HeLa cell (Zhu et al., 2018a), which is estimated to have a volume of a few picoliters (Zhao et al., 2008; Park et al., 2008). We analyzed single cells from E15 chicken utricle; by using peeled epithelia for cell dissociation, we limited the cell types analyzed to hair cells and supporting cells (Herget et al., 2013), each of which is smaller than a HeLa cell. To distinguish the two cell types, we labeled utricle cells with FM1-43, which labels hair cells more strongly than supporting cells (Herget et al., 2013; Ellwanger et al., 2018). After cell dissociation, we collected single cells and pools of 3, 5, and 20 cells in nanowells using fluorescence-activated cell sorting (FACS; Figures 1A and Figure 1—Figure Supplement 1). As expected, collected cells with high levels of FM1-43 (FM1-43_high_) had hair bundles and elongated cell bodies (Figure 1B), both of which are characteristic of hair cells; FM1-43_low_ cells—mostly supporting cells—were round after FACS (Figure 1C). We measured cell volumes after cell sorting. FM1-43_high_ cells averaged 1.01 ± 0.20 picoliters (mean ± SD; *N*=9), while FM1-43_low_ cells averaged 0.67 ± 0.15 picoliters (*N*=8). For comparison, using the same method, we measured the volume of HeLa cells to be 4.9 ± 1.1 picoliters (*N*=5).

**Figure 1.**
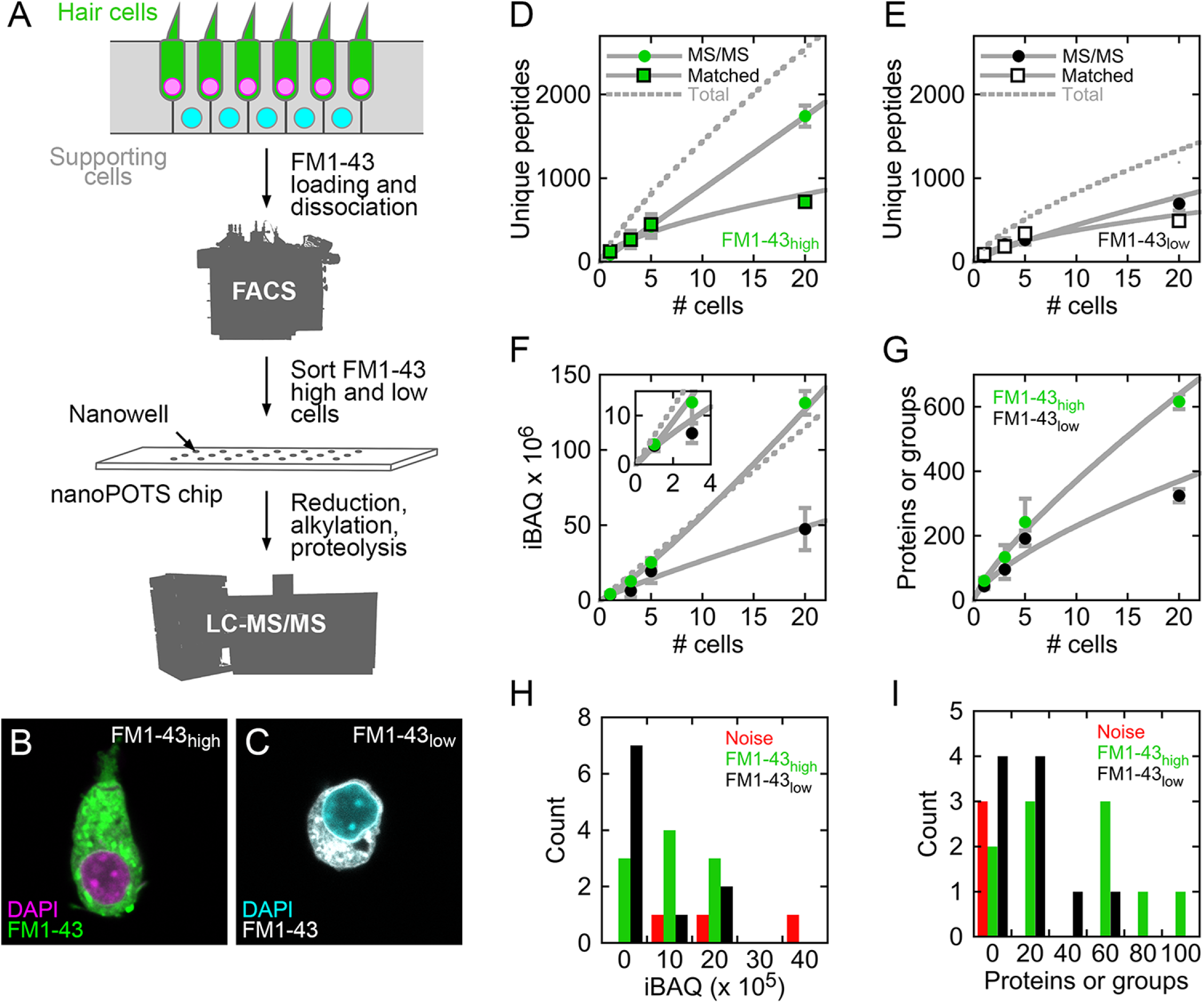
Mass spectrometry of single cells and small cell pools from E15 chick utricle. ***A***, Experimental design. The E15 chick utricle’s sensory epithelium consists of sensory hair cells and supporting cells, which are also progenitor cells. FM1-43 labels hair cells more strongly than supporting cells. The dissociated cells were sorted by FACS and deposited into single nanowells in nanoPOTS chips, where sample processing was carried out without transfer. Samples were loaded into glass microcapillaries and were analyzed by mass spectrometry. LC-MS/MS, liquid chromatography-tandem mass spectrometry. ***B***, FACS-sorted FM1-43_high_ cell with typical hair cell cytomorphology, including apical hair bundle. ***C***, FACS-sorted FM1-43_low_ cell shows rounded cytomorphology after dissociation. For B and C, FM1-43 dye was added again to cells after sorting in order to visualize cell shape; dye intensity therefore is not representative of the signal used for sorting hair cells from supporting cells. ***D-E***, Relationship between number of cells and unique peptides. Peptides directly identified by MS2 (peptide fragmentation) spectrum matching are shown by circles and those indirectly identified by Match Between Runs by squares; data are separately plotted for FM1-43_high_ (D) and FM1-43_low_ (E). Gray solid lines are power fits to data through (0,0); gray dashed line is fit to sum of the MS/MS and Matched data. ***F***, Relationship between number of cells and total iBAQ. Gray solid line is power fit through (0,0); gray dashed line is linear fit through (0,0). Green, FM1-43_high_; black, FM1-43_low_. Inset shows 1-3 cells only. ***G***, Relationship between number of cells and the total number of proteins or protein groups identified. Gray solid line is power fit through (0,0). Data for D-G were from Experiment 1; mean ± SEM are plotted. ***H***, Distribution of total iBAQ for individual cells (FM1-43_high_, green; FM1-43_low_, black) or for blank wells (red). Count refers to the number of cells in a bin. Note that total iBAQ does not distinguish individual cells from noise. ***I***, Distribution of number of identified proteins or protein groups. This measure distinctly distinguishes individual cells from noise; those FM1-43_high_ or FM1-43_low_ samples with low numbers of identifications likely do not have cells in the nanowells. Data for H-I were from Experiment 2.

We used the nanoPOTS (nanodroplet processing in one-pot for trace samples) approach to carry out all sample processing steps in single nanowells (Zhu et al., 2018b). After protein extraction, reduction, alkylation, and proteolysis, digested peptides were collected and separated using nano-liquid chromatography on a 30-µm-i.d. column (Zhu et al., 2018b) (Figure 1A). The separated peptides were delivered to an Orbitrap Fusion Lumos Tribrid mass spectrometer and were analyzed using data-dependent acquisition. Peptides were identified, quantified, and assembled into proteins with Andromeda and MaxQuant (Cox and Mann, 2008; Cox et al., 2011), using Match Between Runs (Tyanova et al., 2016; Zhu et al., 2018b) to identify unmatched peptides based on their accurate masses and liquid chromatography retention times. To quantify proteins based on molar abundance, we used intensity-based absolute quantification (iBAQ), calculated from the sum of peak intensities of all peptides matching to a specific protein divided by the number of theoretically observable peptides (Schwanhäusser et al., 2011). All mass spectrometry data are deposited at ProteomeXchange, and analyzed data are reported in Figure 1—Source Data 1.

About 200 unique peptides were identified in each single FM1-43_high_ cell; about half were identified by MS/MS scans and half by matching (Figure 1D). The total number of unique peptides increased to ∼2500 in pools of 20 cells, with about 70% identified by MS/MS scans (Figure 1D). Only ∼1200 peptides were identified in pools of 20 FM1-43_low_ cells, with about 60% identified by MS/MS scans (Figure 1E). Total iBAQ rose nonlinearly with the number of cells (Figure 1F), suggesting that some protein was lost to surface adsorption; while small relative to typical sample wells, the volume of the nanowell is still 50,000-fold larger than the volume of a utricle cell. Because sample processing occurred in a protected nanowell environment using robotic liquid handling, the total iBAQ attributed to keratins (e.g., human skin contamination) was only ∼0.1% of the total, far less than >50% occurring in some mass-spectrometry experiments with small amounts of protein. The number of proteins or protein groups identified increased from ∼60 for FM1-43_high_ single cells to nearly 600 for pools of 20 cells (Figure 1G); fewer proteins were identified in supporting cells, likely because of their smaller volume.

Comparison of single FM1-43_high_ and FM1-43_low_ cells to wells with collection triggered to noise allowed us to confirm the presence of single cells, even without visual inspection of the wells. Total iBAQ did not accurately indicate which wells contained single cells (Figure 1H), presumably because the total signal can be dominated by incorrect assignment of contaminant signals to proteins. By contrast, the number of proteins or protein groups identified distinguished most FM1-43_high_ or FM1-43_low_ samples from noise (Figure 1I); the samples with low numbers of identifications could represent sorting events where cells missed their target nanowell.

To examine the composition of FM1-43_high_ and FM1-43_low_ samples, we used relative iBAQ (riBAQ) for quantitation (Shin et al., 2013; Krey et al., 2014) and displayed the 60 most abundant proteins of the 20-cell samples (Figure 2A). All identified proteins from the 20-cell samples are displayed in Figure 2—Figure Supplement 1, and all identified proteins from the single-cell samples are displayed in Figure 2—Figure Supplements 2 and 3. We used a volcano plot analysis to show those proteins that had statistically significant enrichment in the 20-cell FM1-43_high_ and FM1-43_low_ samples (Figure 2B). Well-known hair-cell proteins were enriched significantly in FM1-43_high_ cells, including the mobile Ca^2+^ buffers OCM and CALB2, as well as the molecular motor MYO6. Several proteins were highly enriched in the FM1-43_low_ samples, including TMSB4X, STMN2, SH3BGRL, and MARCKS.

**Figure 2.**
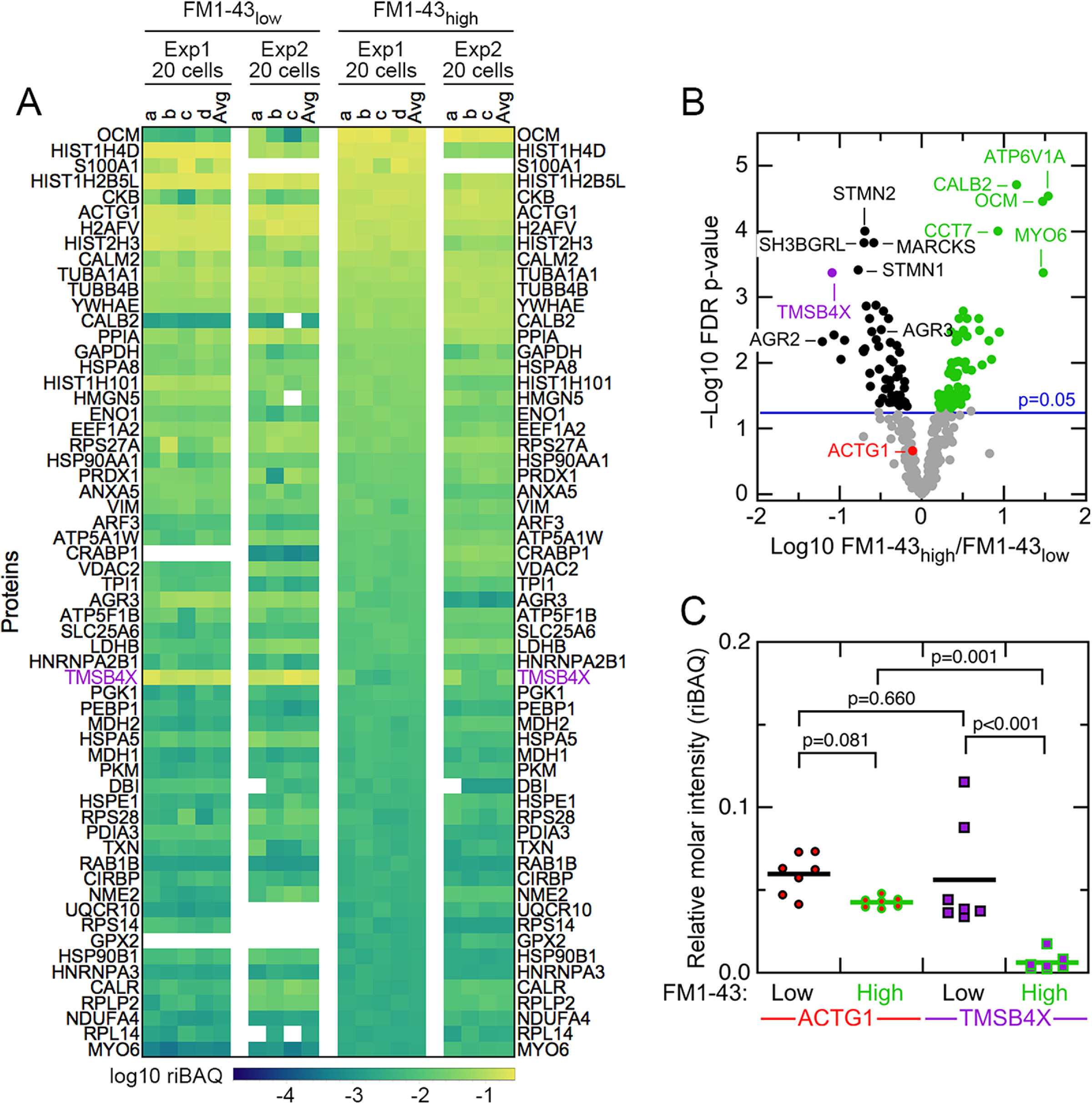
Abundant proteins in small pools of isolated E15 chick utricle cells. ***A***, Heat map showing top 60 proteins or protein groups in samples of 20 cells, sorted by the average of the 20-cell FM1-43_high_ samples. FM1-43_low_ and FM1-43_high_ samples from Experiment 1 (Exp1) and Experiment 2 (Exp2) are both displayed, as are the averages of the individual samples. TMSB4X is called out with magenta type. Scale on bottom indicates relationship between riBAQ and color. ***B***, Volcano plot showing relationship between FM1-43_high_/FM1-43_low_ enrichment (x-axis) and false discovery rate (FDR)-adjusted p-value (y-axis). Proteins that are significantly enriched are labeled with green (FM1-43_high_>FM1-43_low_) or black (FM1-43_low_>FM1-43_high_). ***C***, ACTG1 and TMBS4X quantitation. Relative molar fraction (riBAQ) quantitation of ACTG1 (circle, red fill) and TMSB4X (square, magenta fill) expression in FM1-43_high_ cells (green outline) or FM1-43_low_ cells (black outline). Samples 20 cells are plotted; lines indicate mean expression level for the group. Statistical significance is indicated.

### Characterization of TMSB4X and monomeric actin in chick utricle

The proteomics experiments also revealed several abundant proteins that had not been previously found to be hair-cell specific, including GSTO1, GPX2, CRABP1, and AK1; TMSB4X and AGR3 were examples of proteins that were much more abundant in supporting cells (Figure 1—Source Data 1; Figure 2A-B). We examined several of these proteins in E15 chick utricles using immunocytochemistry. Antibody labeling for AGR3 and the hair-cell marker OTOF labeling did not overlap, and the elongated cell bodies labeled for AGR3 indicated that it marked supporting cells (Figures 3A-C and Figure 3—Figure Supplement 1). By contrast, CRABP1 was specific for hair cells, seen by the overlap with OTOF (Figures 3D-F and Figure 3—Figure Supplement 2).

**Figure 3.**
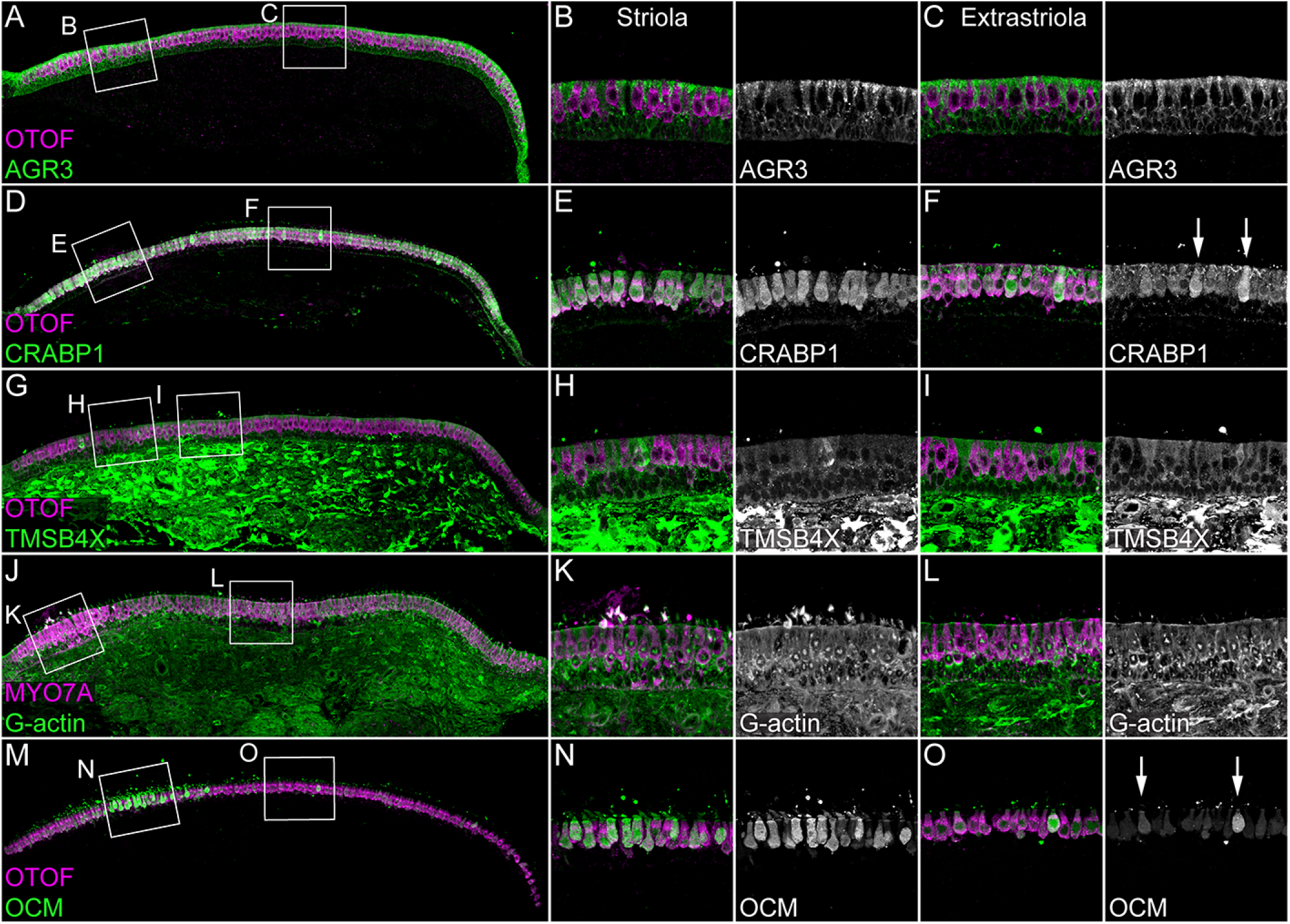
Immunolocalization of proteins enriched in hair cells or supporting cells of E15 chick utricle. Confocal z-stacks of vibratome cross-sections of the whole utricle were imaged with the tiling and stitching function in Zeiss ZEN. Confocal z-stacks for magnified extrastriolar and striolar regions were collected separately. A subset of the z-stacks series was used for the maximum intensity projection to preserve single-cell resolution. Hair cells are labeled with antibodies against OTOF or MYO7A (each in magenta). ***A-C***, AGR3 (green) was detected in extrastriolar and striolar supporting cells. Panel full widths: A, 913 µm; B-C, 125 µm. ***D-F***, CRABP1 (green) is concentrated in hair cells. A few extrastriolar hair cells show very high levels of CRABP1 (arrows). Panel full widths: D, 1038 µm; E-F, 125 µm. ***G-I***, TMSB4X (green) immunoreactivity was intense in cells of the mesenchymal stromal cell layer. TMSB4X was detectable at moderate levels in extrastriolar and striolar supporting cells and at low levels in hair cells located in extrastriolar and striolar regions. Panel full widths: G, 934 µm; H-I, 125 µm. ***J-L***, G-actin (JLA20 antibody, green) is expressed at equal levels in hair cells, supporting cells and mesenchymal stromal cells. Panel full widths: J, 839 µm; K-L, 125 µm. ***M-O***, OCM was detectable at high levels in striolar hair cells and at low levels in extrastriolar hair cells. A few extrastriolar hair cells display high levels of OCM (arrows). Panel full widths: M, 946 µm; N-O, 125 µm. Expanded images of all individual channels (transmitted light, nuclei, F-actin, hair cells, and specific antibody) are shown in Figure 3—Figure Supplements 1-5.

The thymosin-beta family of proteins, which includes TMSB4X, are actin monomer binding proteins that sequester substantial fractions of actin in many cell types (Nachmias, 1993; Sun et al., 1995). Five TMSB4X peptides were identified by mass spectrometry, which covered 75% of the ∼5 kD protein; one of the peptides was shared by TMSB15B, another member of the family (Figure 1—Figure Supplement 2). Analysis of transcript expression in mouse inner ear using gEAR (https://gear.igs.umaryland.edu) indicated that *Tmsb4x* expression was considerably higher than that of another paralog, *Tmsb10*, and much higher than the two *Tmsb15* isoforms, justifying our focus on TMSB4X.

To localize TMSB4X in the E15 chick utricle, we used an antibody that has been validated previously with knock-down experiments against mouse TMSB4X (Zhou et al., 2013; Li et al., 2018); chicken TMSB4X differs from mouse and human TMSB4X by only two serine-to-threonine substitutions out of 44 total amino acids. TMSB4X immunoreactivity was cytoplasmic and strong in supporting cells, but was substantially reduced in hair cells (Figures 3G-I and Figure 3—Figure Supplement 3), which was consistent with the mass-spectrometry results. Because TMSB4X maintains actin in a monomeric form (G-actin), probes for G-actin like the JLA20 antibody (Lin, 1981) provide another way of localizing the pool of unpolymerized actin. JLA20 immunoreactivity was similar to that of TMSB4X, although it was less enriched in supporting cells than was TMSB4X (Figures 3J-L and Figure 3—Figure Supplement 4).

The concentration of TMSB4X relative to total actin indicates how much free actin is available for assembling filamentous structures like stereocilia. Analyzing the 20-cell samples, we found that the ACTG1 protein group—total actin—accounted for a relative molar fraction (riBAQ) of 0.043 ± 0.001 (mean ± SEM) in FM1-43_high_ cells and 0.060 ± 0.005 in FM1-43_low_ cells (Figure 2C). A mixed-effects model accounting for intra-sample correlations indicated that these concentrations differed significantly, albeit only at an alpha level of 0.05 (summary statistics with confidence intervals are reported in Table 1). While TMSB4X accounted for a relative molar fraction of only 0.006 ± 0.002 in FM1-43_high_ cells, it was 0.056 ± 0.012 in FM1-43_low_ cells, ten-fold higher (Figure 2C) and significantly different (p<0.001). Critically, the concentration of hair-cell TMSB4X differed significantly from that of hair-cell actin (p=0.001), while the concentration of supporting cell TMSB4X did not differ from that of supporting cell actin (p=0.660). Because TMSBX and actin interact with a 1:1 stoichiometry (Goldschmidt-Clermont et al., 1992), and no other actin-binding proteins are detected at similar high levels, our quantitation suggests that TMSB4X is capable of binding most actin monomers in supporting cells.

**Table 1.**
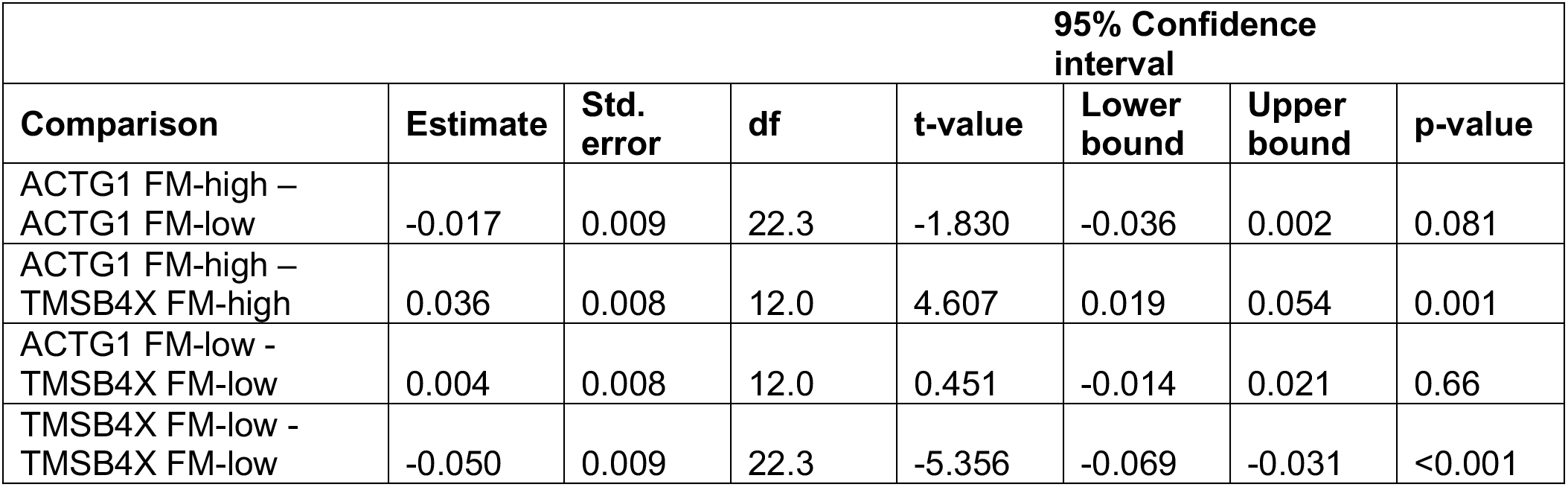
Summary statistics with confidence intervals for Fig. 2C.

In wholemount preparations, we counted 72 ± 8 stereocilia per utricle hair cell (mean ± SD; *N*=26 from striolar and extrastriolar regions). The actin quantitation suggested that each E15 hair cell contains ∼15,000,000 actin molecules (G- and F-actin combined). If nearly all actin is in stereocilia (Tilney and Tilney, 1988), then each stereocilium would contain ∼200,000 actin molecules. While fewer than the 400,000 molecules estimated per E20 chick stereocilium (Shin et al., 2013), the value is consistent with the relative immaturity of E15 cells.

### Developmental trajectory analysis using single-cell proteomics

While our results showed that the expression profile of pooled-cell samples distinguished between supporting cells and hair cells, a single contaminating cell in a pool could distort the pool’s expression pattern. We therefore examined whether we could achieve similar discrimination based on the 30 single-cell profiles, despite the low numbers of identifications in each cell. We used 75 proteins or protein groups that were detected in at least five cells, and normalized and batch-corrected (Johnson et al., 2007; Büttner et al., 2019; Luecken and Theis, 2019) data from individual cells. The resulting expression values, referred to as log2 normalized iBAQ (niBAQ) units, comprised an expression matrix having similar characteristics to single-cell transcriptomics data. Non-detects accounted for 62% of the values, while for 83% of identifications the detected values followed a log-normal distribution (Shapiro-Wilk normality test p>0.05).

Because methods developed for single-cell transcript analysis (Luecken and Theis, 2019) should be suitable for dissection of the single-cell proteomics results, we applied CellTrails, which we previously used to uncover the branching trajectory from progenitors to hair cells in the chicken utricle using transcript data (Ellwanger et al., 2018). To interpret the latent structure in the single-cell mass spectrometry data, its lower-dimensional manifold was investigated using CellTrails’ robust nonlinear spectral embedding on the submatrix of the 37 highest variable identifications (Figure 4A). Appropriately, the cells distinctly segregated according to their FM1-43 uptake (Figure 4B). We noted that the protein pattern of three cells classified as FM1-43_high_ appeared to match better to the FM1-43_low_ (supporting cell) pattern. Similarly, two FM1-43_low_ cells were embedded in the neighborhood of cells with a high FM1-43 uptake (hair cells). While FM1-43 is useful for labeling hair cells, transcript analysis showed that FM1-43 levels are not a perfect proxy for hair-cell maturity (Ellwanger et al., 2018); for example, hair cells with damaged mechanotransduction will not load with the dye and such cells would be classified as FM1-43_low_. Alternatively, cells with relatively low FM1-43 could be transitional cells between progenitors and mature hair cells (Ellwanger et al., 2018). We therefore surmised that we could elicit a developmental trajectory from the single-cell protein expression patterns. The chronological ordering of the cells was learned in the lower-dimensional manifold and a pseudotime value was assigned to each cell (Figure 4C).

**Figure 4.**
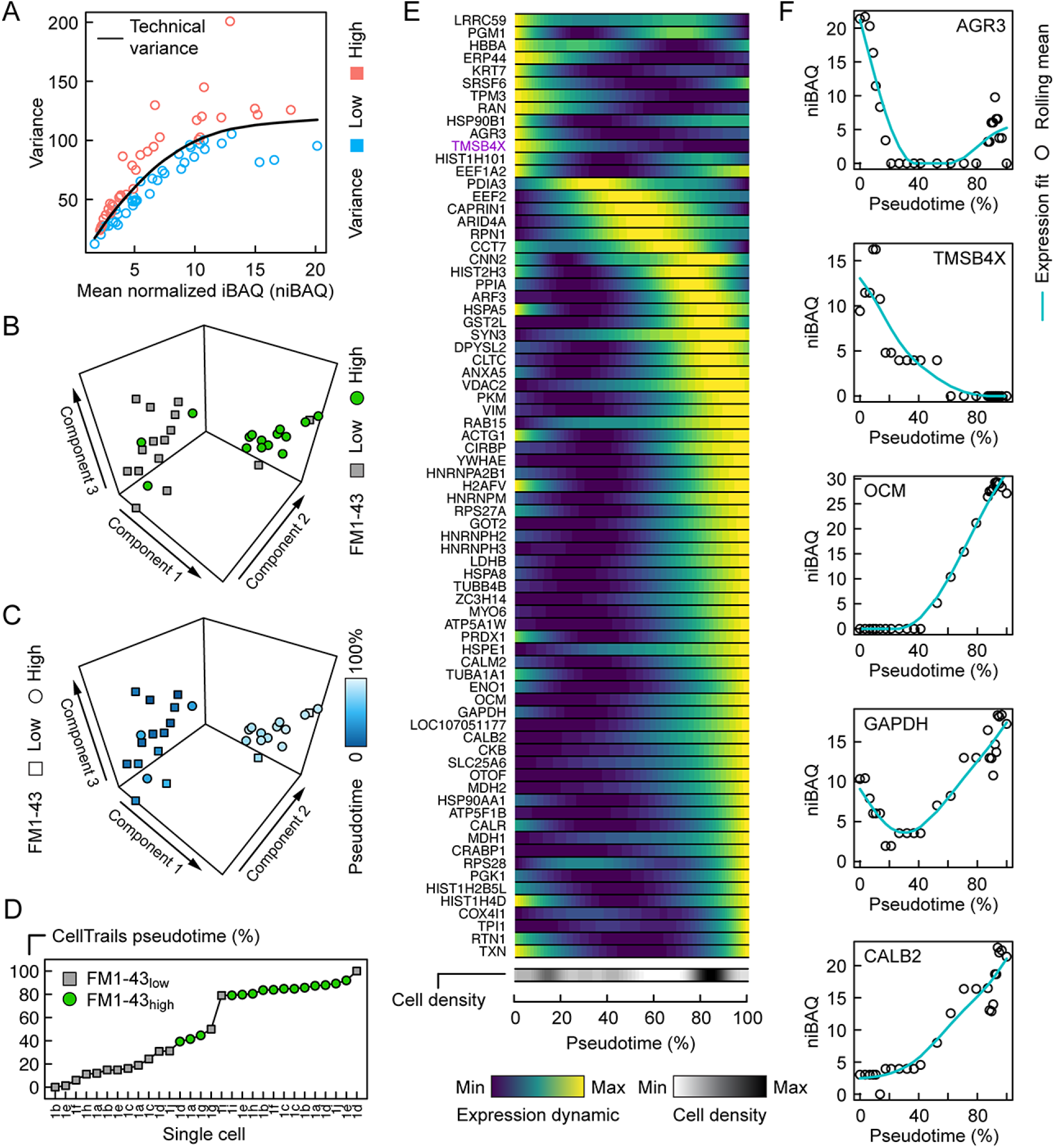
Pseudotemporal ordering of single utricle cells based on proteomics measurements. ***A***, Relationship between variance and mean expression, distinguishes proteins or protein groups with low variance (blue) or high variance (salmon). ***B***, First three components of CellTrails’ spectral embedding, with FM1-43_low_ (square, gray) and FM1-43_high_ (circle, green) cells indicated. ***C***, First three components of CellTrails’ spectral embedding with cells colorized by the inferred pseudotime. ***D***, Chronological ordering of single cells as a function of pseudotime shows that cell ordering correlates with the FM1-43 uptake gradient. ***E***, Scaled expression dynamics over pseudotime for all analyzed proteins or protein groups. A cubic smoothing spline with four degrees of freedom was fit on the rolling mean for each protein. Cell density bar underneath the heat map shows the density of cells along the pseudotime axis. Heat map and cell density scale is shown below. ***F***, Absolute expression dynamics of log2 niBAQ expression levels as a function of pseudotime for various proteins. Blue line is expression fit; circle is the rolling mean for each protein.

The 75 proteins sufficiently detected on single-cell level are all relatively highly expressed and largely do not include those expected to distinguish different classes of hair cells (Ellwanger et al., 2018). Moreover, we expect that our sample is dominated by type II hair cells, especially those from extrastriola regions, as they are much more numerous than type I hair cells (Ellwanger et al., 2018). Because of the gating strategy used (Figure 1—Figure Supplement 1), we primarily sampled cells from either end of the developmental trajectory, which was apparent from the gap approximately at the midpoint of the pseudotime axis (Figure 4D). Nevertheless, we sampled sufficient numbers of differentiating cells to establish a developmental trajectory.

We examined protein expression dynamics as a function of developmental pseudotime (Figure 4E). Protein expression changed systematically along the developmental pseudotime axis with the expected trends: proteins enriched in supporting cells, including AGR3 and TMSB4X, decreased in expression along the pseudotime axis (Figure 4E-F). By contrast, proteins known to be enriched in hair cells, including OCM, CALB2, MYO6, CKB, and GAPDH, all increased as pseudotime progressed (Figure 4E-F).

### Transcriptomic confirmation of TMSB4X enrichment in progenitor cells

We predicted that the decrease in TMSB4X as hair cells mature arose from downregulation of *TMSB4X* transcript expression during differentiation of hair cells. We therefore used transcriptomic profiling of single cells isolated from E15 chick utricle to examine gene expression during the bifurcating trajectory that describes the development of progenitor cells to mature striolar and extrastriolar hair cells (Ellwanger et al., 2018). We carried out scRNA-seq transcriptomic profiling using the Smart-seq protocol (Picelli et al., 2014) on 384 FACS-sorted E15 chick utricle hair cells. To provide maximum correlation of *TMSB4X* expression changes with chicken utricle hair cell maturation, we reconstructed the trajectory in similar fashion as previously described (Ellwanger et al., 2018), carrying out the analysis with 182 assay genes already including *GSTO1* and *CRABP1* from that previous study, supplemented with *TMSB4X*, *AGR3*, *GPX2* and *AK1*. Nine cellular subgroups emerged, each of which was distinguished by distinct marker gene sets (Figure 5A). Based on their expression profiles, for example the lack of *TECTA* and especially high levels of *TMSB4X* (Figure 5A; see also Figure 3), two subgroups (S8 and S9) appeared to be stromal cells; to focus on the developmental progression of progenitor (supporting) cells to hair cells, we removed S8 and S9 for subsequent analysis.

**Figure 5.**
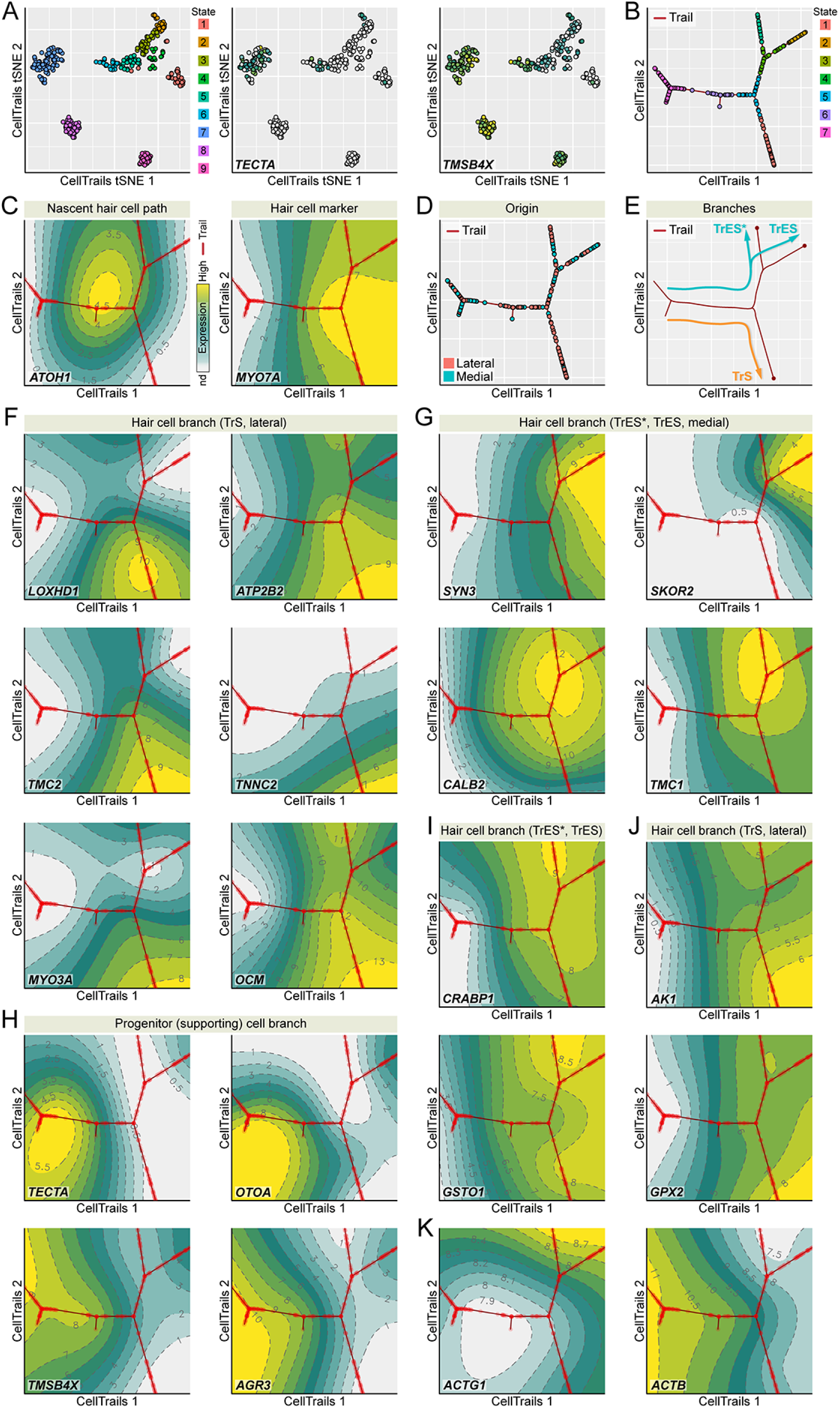
Developmental trajectory identified from single utricle cell RNA-seq measurements. ***A***, CellTrails identifies nine distinct states (cellular subgroups). Shown are 328 points representing single utricle cells projected into two-dimensional space using CellTrails t-distributed stochastic neighbor embedding (tSNE). Cells are colored by state affiliation. Two states (S8-9) were classified as stromal cells and excluded from this study based on the lack of *TECTA* expression and the high levels of *TMSB4X* expression. ***B***, CellTrails trail map of 254 single chicken utricle cells reveals a bifurcating trajectory. ***C***, *ATOH1* peaks before the main bifurcation of the right major branches. CellTrails map shows that *MYO7A*-expressing hair cells are located downstream of the *ATOH1* peak on the right half of the trajectory map. ***D***, Projection of medial and lateral cell origin metadata into the trail map. Cells from the lateral side accumulate along the lower-right trajectory whereas cells from both halves are located along the upper-right trajectory. ***E***, Predicted developmental extrastriolar (TrES*, TrES), and striolar (TrS) hair cell trajectories. ***F***, *LOXHD1, ATP2B2*, *TMC2*, *TNNC2*, *MYO3A,* and *OCM* expression levels are associated with the lower-right lateral striolar (TrS) branch. ***G***, CellTrails maps showing high expression of *SYN3*, *SKOR2*, *CALB2,* and *TMC1* along the upper-left medial extrastriolar (TrES*, TrES) branch. ***H***, Expression of supporting cell marker genes *TECTA*, *OTOA*, *TMSB4X* and *AGR3* defines the location of the progenitor (supporting) cell population along the left major branch. ***I***, Expression of *CRABP1* and *GSTO* is associated with the extrastriolar trajectory* (TrES*, see also Figure 3G, high *CRABP1* expression extrastriolar hair cells). ***J***, *AK1* and *GPX2* are enriched in the striolar trajectory (TrS). ***K***, *ACTB* expression decreases and *ACTG1* expression increases while hair cells develop.

We mapped the remaining 254 individual cells of subgroups S1-S7 along developmental trajectories, plotting CellTrails maps (Ellwanger et al., 2018) to demonstrate the branching nature of the trajectory (Figure 5B). Our assay was biased for hair-bundle genes, and at least half of the cells isolated were hair cells, so it is unsurprising that the final trajectory revealed not only the transition from progenitor (supporting) cells to hair cells, but also further developmental branching. One major branch was supporting cells, as these cells expressed markers like *TECTA* and *OTOA* (Figure 5H; Figure 5—Source Data 1). The other two major branches were hair cells, as they occurred after the peak of *ATOH1* expression and also showed high levels of hair-cell markers like *MYO7A* (Figure 5C; Figure 5—Source Data 1).

Enrichment of *LOXHD1, ATP2B2*, *TMC2*, *TNNC2*, *MYO3A* and *OCM* indicated that the branch projecting down and to the right was from striolar cells (Figure 5F; Figure 5—Source Data 1), which we named TrS (“trail striola”) as previously defined(Ellwanger et al., 2018). Similarly, enrichment of *SYN3*, *SKOR2*, *CALB2*, and *TMC1* indicated the branch projecting up and to the right is from extrastriolar cells, which we named TrES. TrES branched again, and relative enrichment of *ATP2B2* (Figure 5F; Figure 5—Source Data 1) and reduced expression of *SKOR2* (Figure 5G; Figure 5—Source Data 1) indicated that the left-hand branch was equivalent to the novel hair cell type TrES* found in our previous study (Ellwanger et al., 2018).

To confirm the spatial identity of the two major hair cell branches, all experiments were carried out with E15 chicken utricles split apart into lateral halves, which contain striolar and extrastriolar cells, and medial halves, which contain only extrastriolar cells. The lower-right branch was populated nearly entirely by lateral cells, which confirms that it represents striolar hair cells (Figure 5D-E). We conclude that the scRNA-seq experiment accurately replicated our previous experiment using a multiplex RT-qPCR approach (Ellwanger et al., 2018).

We next examined the genes highlighted in the proteomics experiments, including *TMSB4X*, *AGR3*, *GSTO1*, *GPX2*, *CRABP1*, and *AK1*. As predicted from the proteomics and localization experiments, the CellTrails analysis showed that *TMSB4X* and *AGR3* were specific to progenitor (supporting) cells (Figure 5H), while *GSTO1*, *GPX2*, *CRABP1*, and *AK1* were specific to hair cells (Figure 5I). The CellTrails maps indicated that *GPX2* and *AK1* were concentrated in striolar hair cells, while *CRABP1* was enriched in extrastriolar cells, particularly TrES* (Figure 5I). Cells observed with high levels of CRABP1 in immunocytochemistry experiments could be the TrES* cells (Figure 3G,I). *GSTO1* was expressed at similar levels in both hair cell types. Available antibodies against GPX2, AK1, and GSTO1 were insufficiently reliable to check their hair-cell specificity. Examining databases in gEAR, however, we noted that *Gpx2* and *Ak1* are predicted to be substantially enriched in hair cells as compared to non-hair cells in mouse utricle; by contrast, *Gsto1* is expressed at higher levels in mouse utricle non-hair cells than in hair cells.

The scRNA-seq results corroborated the expression dynamics of TMSB4X on transcriptional level. High in progenitor cells, its transcriptional activity decreased substantially during hair cell differentiation (Figure 5H), and was nearly undetectable in striolar hair cells. Interestingly, *TMSB4X* was expressed at detectable levels, albeit relatively low, in cells along TrES as compared to TrES* (Figure 5H).

We also noted a striking decrease in *ACTB* expression as hair cells differentiated; the CellTrails maps suggested that *ACTB* was >10-fold higher in progenitor cells than in hair cells (Figure 5K; Figure 5—Source Data 1). *ACTG1* increased modestly in expression, especially in TrES cells, but overall was present at lower levels than *ACTB* (Figure 5K; Figure 5—Source Data 1). Similar trends for these actin isoforms were also seen in our previous data (Ellwanger et al., 2018). ACTB and ACTG1 differ by only four amino acids, however, and we only detected one of the peptides that distinguish the isoforms (Ac-EEEIAALVIDNGSGMCK from ACTG1) in single mass spectrometry run. We were therefore unable to accurately measure the relative abundance of the two actin isoforms in our protein mass spectrometry experiments.

## Discussion

Global analysis of proteins from single cells has previously been thwarted by nonspecific adsorption of proteins to surfaces and the lack of a scheme to amplify proteins (Couvillion et al., 2019). Recent development of nanowell sample processing with nanoPOTS, coupled with extremely sensitive mass spectrometers, permits detection of abundant proteins of even very small cells (<1 picoliter). We exploited this method to characterize the abundant proteins of FACS-sorted supporting cells, the hair cell progenitors, and hair cells from embryonic chick utricles. Remarkably, we were able to use the mass spectrometry data from single utricle cells to reconstruct a developmental trajectory from protein-expression values alone.

In addition, we identified several proteins not previously highlighted as specific for hair cells (CRABP1, GSTO1, GPX2, AK1) or for supporting cells (AGR3, TMSB4X). TMSB4X was present at nearly equimolar levels with respect to actin in supporting cells, indicating that most actin is sequestered in those cells. By contrast, in hair cells, TMSB4X was only one-tenth as abundant as actin. This developmental change was characterized in more depth using single-cell RNA sequencing, which showed that the drop in *TMSB4X* was greater in extrastriolar hair cells than in striolar hair cells. Together, these data strongly suggest that downregulation of TMSB4X allows differentiating hair cells to construct their hair bundles with newly available actin monomers.

### Single-cell proteomics detection

While fewer proteins were identified from single hair cells than were identified in a recent report using the same technique with single HeLa cells and primary lung cells (Zhu et al., 2018a), hair cells have a volume of ∼1 picoliter, while HeLa cells are five times larger in volume. Given this difference, detection of fewer proteins in hair cells than in HeLa cells was not surprising.

While the nanoPOTS approach is useful for characterizing abundant proteins in small cells or many proteins in larger cells, without further increases in sensitivity, the small number of proteins detected in single hair cells will prevent characterization of low-abundance proteins or deeply categorizing developmental pathways using protein expression. Because the relationship between cell number and total protein signal intensity was nonlinear, especially for 1-3 cell samples (Figure 1F), we concluded that we lost significant amounts of protein to adsorption to the nanowells. These results indicate that further improvement of protein recovery is critical to increase proteome coverage and quantification performance of single-cell proteomics technology employing nanoPOTS. Fabricating nanowells with smaller dimensions would be straightforward; however, dispensing single cells by FACS to yet-smaller wells would be difficult to carry out reproducibly. In addition, the nanowell surfaces could be coated chemically with antifouling materials such as polyethylene glycol or poly(2-methyl-2-oxazoline) polymers (Weydert et al., 2017). An alternative strategy for single-cell protein analysis uses TMT multiplex labeling with one channel utilized by a sample of several hundred carrier cells, which will reduce the relative error due to protein loss(Budnik et al., 2018). Coupling of TMT multiplex labeling approach with nanoPOTS could significantly increase proteome coverage and analysis throughput of single cell proteomics.

We detected much higher levels of small proteins (<20 kD) here than in previous experiments using the same tissue (Shin et al., 2007; Shin et al., 2010; Shin et al., 2013; Herget et al., 2013; Wilmarth et al., 2015). We attribute this improved detection to the processing of samples in nanoPOTS nanowell without transfer steps; other sample preparation methods included steps (e.g., SDS-PAGE gel separation or filter washes) that facilitated loss of small proteins (Krey et al., 2016). Our new data indicate that on a molar basis, OCM (Figure 2A, Fig 3M-O), which was formerly known as parvalbumin 3 in the chick (Heller et al., 2002), is the most abundant protein in E15 chick hair cells, accounting for >10% of the total number of protein molecules.

### Single-cell proteomics trajectory analysis

We report here the first example of the use of single-cell protein mass spectrometry data to construct a developmental trajectory, in this case from progenitors to hair cells. While limited by having only 30 cells for this analysis, CellTrails nevertheless was able to generate a developmental trajectory that discriminated distinct expression patterns of transitioning cells. The proposed trajectory for the single cells studied here accounted for protein expression levels at the endpoints, which represented supporting cells and hair cells; moreover, expression levels of many of the endpoint-selective proteins systematically increased or decreased across the trajectory, as expected. Although we did not detect the branching trajectories of hair cells along the TrS, TrES, and TrES* trails that were seen using multiplex RT-qPCR approach (Ellwanger et al., 2018) or scRNA-seq (Figure 5), the small number of cells analyzed for protein-based trajectories likely precluded their detection.

Although application of trajectory-analysis methods to single-cell proteomics is very much in its infancy, we show here that CellTrails is suitable for this purpose. The analysis was limited by sensitivity, as proteins detected by mass spectrometry were limited to those expressed at relatively high levels. Improvements of the nanoPOTS method that lead to increased sensitivity and reproducibility will enhance future protein-based trajectory analyses. Nevertheless, while robotic manipulation allows for increased sample-preparation output, the number of cells analyzed presently must remain low because of slow throughput of the mass spectrometry steps. Single-cell RNA-seq approaches are likely to continue to offer much higher throughput and depth for the foreseeable future. That said, analysis of developmental pathways using single-cell proteomics allows the identification of key proteins that change in protein expression level without alternations in transcript levels. Moreover, future single-cell proteomics approaches will allow analysis of posttranslational modifications like phosphorylation, which will expand our ability to probe developmental cascades.

Interestingly, GAPDH was found to increase during hair cell maturation (Figure 4E-F), while its mRNA was reported to remain at a constant level (Avenarius et al., 2014; Ellwanger et al., 2018); this discrepancy suggests that GAPDH undergoes post-transcriptional regulation, either from increased translation efficacy or by protein stabilization following translation. Because GAPDH concentrates in stereocilia (Shin et al., 2007; Shin et al., 2013), protein stabilization there is the most likely explanation for the increasing GAPDH levels during hair-cell differentiation. This observation highlights the power of the protein-based trajectory analysis; because of the poor correlation between transcript levels and protein levels (Liu et al., 2016), understanding how proteins change during a developmental process will require their direct measurement, not inferring their levels as is done with scRNA-seq experiments.

### Actin expression dynamics

During formation of hair cells, we observed a substantial downregulation of *ACTB* during differentiation to hair cells, with the largest decrease in extrastriolar cells; this decrease was partially compensated by an increase in *ACTG1*, especially in extrastriolar cells. If scRNA-seq provides accurate relative measurements of transcript levels, the sum of *ACTB* and *ACTG1* transcripts in E15 hair cells was one-quarter that in supporting cells. Total actin protein levels were similar in hair cells and supporting cells, however, suggesting that actin synthesis may be reduced during differentiation of hair cells.

In chicken auditory hair cells, ACTB is found nearly exclusively in stereocilia F-actin cores, while the more abundant ACTG1 is distributed to all F-actin assemblies (Höfer et al., 1997). Nevertheless, *Actb* or *Actg1* knockout mice have normal stereocilia development, suggesting that the two isoforms are interchangeable in mice, at least initially (Belyantseva et al., 2009; Perrin et al., 2010). Phenotypes of aging hair cells from the two mutants differ, however, and these differences arise from the ACTB and ACTG1 proteins themselves, not expression dynamics (Perrin et al., 2010; Patrinostro et al., 2018).

### TMSB4X dynamics and assembly of stereocilia

Studied thoroughly in other cell types, members of the beta-thymosin family have not been previously characterized in the inner ear. As in other cell types (Weber et al., 1992; Carlier et al., 1993), our quantitative proteomics experiments indicate TMSB4X is highly expressed in the inner-ear sensory epithelium; after histones and actin, TMSB4X is the fifth-most-abundant protein in supporting cells (Figure 1—Source Data 1).

While TMSB4X is present at high levels in supporting cells, presumably sequestering actin monomers, it drops substantially in concentration after cells differentiate to hair cells. If total protein in each cell type is the typical ∼250 mg/ml (Fulton, 1982; Srivastava and Bernhard, 1986; Brown, 1991) at an average 56,000 g/mol (Shin et al., 2013), then supporting cells have 0.25 mM TMSB4X and 0.27 mM total actin. Given the 1:1 binding stoichiometry for TMSB4X and G-actin (Goldschmidt-Clermont et al., 1992), only 0.02 mM of actin should be uncomplexed. By contrast, while hair cells have 0.19 mM total actin, their concentration of TMSB4X is only 0.03 mM, leaving 0.16 mM actin uncomplexed and available for building stereocilia.

The changes in TMSB4X expression during development were confirmed and elaborated on by RNA-seq transcriptomic profiling. *TMSB4X* was at high levels in supporting cells and decreased as the cells proceeded towards differentiation as hair cells (Figure 5H; Figure 5—Source Data 1). Striolar hair cells in particular lost nearly all of their *TMSB4X* transcripts. Extrastriolar cells were different; after reaching a nadir, *TMSB4X* increased in TrES cells but remained low in TrES* cells (Figure 5H; Figure 5—Source Data 1).

These results suggest that supporting cells maintain actin in an uncomplexed form, preventing formation of elaborate F-actin arrays (e.g., stereocilia). Upon differentiation to hair cells, however, *TMSB4X* is downregulated, and once the TMSB4X protein has been degraded, actin will be available for stereocilia formation. Thus downregulation of *TMSB4X*, along with the increased transcription of mRNA molecules encoding actin crosslinkers and other F-actin-binding proteins (Ellwanger et al., 2018), is a critical step in formation of the hair cell’s sensory hair bundle.

The reduced expression of actin genes is broadly consistent with Tilney’s suggestion that each hair cell uses an equivalent-sized bolus of actin to build their hair bundles (Tilney and Tilney, 1988). An alternative model dictates that the final amount of actin used in a hair bundle depends on the expression level crosslinkers like PLS1 and ESPN (Höfer et al., 1997; Sekerkova et al., 2011; Krey et al., 2016). A plausible hypothesis incorporating these observations is that ACTB is sequestered with TMSB4X in supporting cells; upon differentiation to hair cells, ACTB is made immediately available for stereocilia elongation by degradation of the actin buffer TMSB4X, while ACTG1 expression is increased to provide actin for other assemblies, including the cuticular plate and circumferential actin belt (Höfer et al., 1997).

## Acknowledgements

We thank Pamela Canaday and Matthew Lewis for their assistance with flow cytometry. We received support from the following OHSU core facilities: FACS from the OHSU Flow Cytometry Shared Resource (supported in part by the OHSU Knight Cancer Institute and the NCI Cancer Center Support Grant P30 CA069533) and confocal microscopy from the OHSU Advanced Light Microscopy Core (P30 NS061800 provided support for imaging). We also received support from the following Stanford core facilities: the Stanford Shared FACS Facility, the Stanford Functional Genomics Facility, and the Otolaryngology Imaging Core. Some of the computing for this project was performed on the Sherlock cluster; we thank the Stanford Research Computing Center for providing computational resources and support. YZ was supported by Earth & Biological Sciences Directorate Mission Seed under the Laboratory Directed Research and Development Program at PNNL, and the Precision Medicine Innovation Co-Laboratory (PMedIC), a joint research collaboration of OHSU and PNNL. A portion of the research at PNNL was performed using EMSL (grid.436923.9), a DOE Office of Science User Facility sponsored by the Office of Biological and Environmental Research. SH was supported by R01 DC015201, by the Hearing Health Foundation’s Hearing Restoration Project, and through the Stanford Initiative to Cure Hearing Loss. RTK was supported by R33 CA225248. PGBG was supported by NIH grant R01 DC011034.

## Author contributions

YZ designed experiments, carried out the single-cell proteomics experiments, and edited the manuscript; MS designed experiments, carried out the single-cell transcriptomics and immunocytochemistry experiments, prepared figures, and edited the manuscript; JK designed experiments, carried out FACS sorting for single-cell proteomics experiments, and edited the manuscript; DE conducted the developmental trajectory analysis using single-cell proteomics data, prepared figures, and edited the manuscript; DC carried out statistical analyses; SH designed experiments, analyzed data, and edited the manuscript; RK designed experiments, analyzed data, and edited the manuscript; PBG designed experiments, analyzed data, prepared figures, and wrote the manuscript.

## Materials and Methods

### Single-cell collection in nanowells and sample preparation for proteomics

Single cells were collected from utricles of E15 chick embryos using methods previously described (Ellwanger et al., 2018). Utricles were incubated with 10 µM FM1-43FX (Thermo Fisher, Waltham, MA) for 20 sec, then treated with thermolysin to remove the sensory epithelium; cells in the epithelium were dissociated by treating with Accutase and mechanical trituration (Ellwanger et al., 2018). Cells were sorted with a BD Influx Instrument (BD Biosciences) set to ‘‘single cell’’ mode and equipped with a 85 µm nozzle. To enable the direct cell sorting into nanowells, a customized template was built with the BD FACS software environment (Sortware) to match the format of nanowell array. SPHERO Drop Delay Calibration Particles (Spherotech, Lake Forest, IL) were used to confirm the drop targeting, as well as the parameter optimization. Prior to sorting onto the final collection chip, a separate preparation of E15 chick cells was used to optimize drop delay settings in order to verify alignment of droplets within the well and cell numbers within each well by visual inspection under an epifluorescence microscope. Debris was removed based on forward scatter (FSC) versus side scatter (SSC) and doublets were excluded based on forward scatter (FSC) versus trigger pulse width. SYTOX Red Dead Cell Stain (Thermo Fisher) was added to cells prior to sorting and was used to identify live and dead cells. Based on our final gating approach, which compared 638-1 (SYTOX Red) versus 488-1 (FM1-43), single SYTOX Red-negative FM1-43_high_ or FM1-43_low_ cells were deposited into individual nanowells.

NanoPOTS chips were fabricated on standard microscopy slides as described (Zhu et al., 2018b; Zhu et al., 2018a). An arrangement of 5 x 13 hydrophilic nanowells was created, each 1 mm diameter with 2.25 mm on-center spacing, and the surrounding chip surface was treated with 2% (v/v) heptadecafluoro-1,1,2,2-tetrahydrodecyl)-dimethylchlorosilane (PFDS) in 2,2,4-trimethylpentane to render it hydrophobic. A glass spacer and cover plate were fabricated for each nanowell chip, allowing the chip to be sealed so that evaporation was minimized during sample incubation. A home-built robotic liquid handling system, capable of subnanoliter dispensing, was used to dispense sample preparation reagents into nanowells (Zhu et al., 2018b; Zhu et al., 2018a). To lyse cells, and to extract and reduce proteins, 100 nl of 0.2% dodecyl β-D-maltoside containing 5 mM DTT in 0.5x PBS and 25 mM ammonium bicarbonate were added to a nanowell containing one or more FACS-sorted samples containing single cells or pools of cells; the chip was then incubated at 70°C for 1 h. Next, proteins were alkylated using 50 nl of 30 mM iodoacetamide in 50 mM ammonium bicarbonate in the dark at 37°C for 30 min. Lys-C and trypsin were added sequentially using 50 nl of 5 ng/μl enzyme solutions in 50 mM ammonium bicarbonate for 4 h and 6 h, respectively. Finally, the peptide sample was acidified with 50 nl of 5% formic acid and then collected into a fused-silica capillary (200 μm i.d., 5 cm long). To maximize sample recovery, each nanowell was re-extracted twice, each with 200 nl of 0.1% formic acid in water. The sample collection capillaries were sealed on both ends with Parafilm and stored at −70°C until use.

Two single-cell proteomics experiments were carried out. For Experiment 1, the FM1-43_high_ samples included five samples with 1 cell, four samples with 3 cells, four samples with 5 cells, and four samples with 20 cells; the FM1-43_low_ samples included five samples with 1 cell, three samples with 3 cells, four samples with 5 cells, and four samples with 20 cells. For Experiment 2, the FM1-43_high_ samples included ten samples with 1 cell and four samples with 20 cells; the FM1-43_low_ samples also included ten samples with 1 cell and four samples with 20 cells.

Images of individual FM1-43_high_ and FM1-43_low_ cells (Figure 1) were acquired by sorting 1000 cells of either cell type into a drop of 4% paraformaldehyde, which had been placed on a Superfrost Plus glass slide (Fisher Scientific). We used FM1-43 to label the isolated cells after fixation; here, the dye’s propensity to insert into the extracellular leaflet of the membrane was used rather than its ability to enter transduction channels. Cells were fixed for 10 min, then a PBS solution containing DAPI and 10 µM FM1-43FX was added to the slide for 10 min. Fixed and labeled cells were washed, then covered with Vectashield mounting medium (Vector Labs) and imaged using a 63x lens on a Zeiss Elyra PS.1 microscope.

### Other quantitation methods

To quantify cell volume, cells were FACS-sorted (for hair cells and supporting cells), fixed, stained with DAPI and phalloidin, and imaged as described above. For each slice of the z-stack, the Threshold and Make Binary tools of Fiji/ImageJ were used to generate a binary stack, which defined the cell perimeter in each z-slice. The Analyze Particles tool was then used to determine the cell area in each slice. The volume for a slice is the product of the single-slice area multiplied by the z-stack interval; all slice volumes were added together to estimate total cell volume.

To count stereocilia per hair bundle, Airyscan z-stack images of phalloidin-stained E15 chicken utricles were obtained using a Zeiss LSM 880 microscope; images were acquired near the base of bundles to ensure that all stereocilia were in each image. Stereocilia were manually counted from single x-y images.

### Data-dependent acquisition mass spectrometry

A capillary solid phase extraction (SPE) column (75 μm i.d., with 3 μm C18 particles of 300 Å pore size; Phenomenex, Torrance, USA) was used for initial sample loading and desalting, and was then connected to a 50 cm, 30 μm i.d. column packed with the same material. Mobile phase was delivered at 60 nl/min with a Dionex UltiMate NCP-3200RS pump system (Thermo Fisher). Peptides were separated with a linear 8-22% Buffer B (0.1% formic acid in acetonitrile) gradient over 60 min, followed by a 10-min increase to 45%. The column was washed with 80% Buffer B for 10 min and then equilibrated with 2% Buffer B for 15 min. An Orbitrap Fusion Lumos Tribrid mass spectrometer (Thermo Fisher) was used for data collection. Peptides were ionized at a spray voltage of 2 kV and ions were collected into an ion transfer capillary set at 150°C. The RF lens was set at 30%. MS1 scans used a 375-1575 mass range, a scan resolution of 120,000, an AGC target of 3 x 10^6^, and a maximum injection time of 246 ms. Precursor ions were selected for MS/MS sequencing if they had charges of +2 to +7 and intensities >8,000; precursors were isolated with an m/z window of 2 and fragmented by high energy dissociation (HCD) set at 30%. Repeat sampling was reduced by using an exclusion duration of 40 s and m/z tolerance of ±10 ppm. MS2 scans were carried out in the Orbitrap with an AGC target of 2 x 10^5^. The maximum injection time and MS2 scan resolution were set as 502 ms and 120,000, respectively.

Andromeda and MaxQuant (version 1.5.3.30) were employed for database searching and label-free protein quantification (Cox and Mann, 2008; Cox et al., 2011). All MS2 spectra were searched against the NCBI Genome Reference Consortium Chicken Build 6a (GRCg6a) database (49,673 protein sequences; released 2018-03-27). The default MaxQuant contaminants file was edited as described (Wilmarth et al., 2015). Carbamidomethylation was selected as fixed modification, and N-terminal protein acetylation and methionine oxidation were set as variable modifications. Peptides must contain >5 amino acids and peptide masses must be <4600 Da; two missed cleavages were allowed for each peptide. Peptides and proteins were each filtered with a false discovery rate (FDR) of 0.01. The Match Between Runs algorithm was used to improve proteome coverage; we used an alignment window of 15 min and a match time window of 0.5 min. iBAQ protein intensities were used for quantification.

### Statistical analysis of proteomics data

For enrichment analysis of FM1-43_high_ (*N*=4 of 20-cell samples) and FM1-43_low_ (*N*=4 of 20-cell samples) cells, the riBAQ data were transformed into log2 scale; linear models with empirical Bayes statistics were fitted to the transformed data using the limma R package (Ritchie et al., 2015) (10.18129/B9.bioc.limma). For the enrichment analysis, we only used the 345 proteins that were measured in at least two replicates in each group. To correct for multiple tests (Benjamini and Hochberg, 1995), the FDR was used to correct two-sided p-values from a moderated t-test (Ritchie et al., 2015); an enrichment of >1.5-fold and a FDR-adjusted p-value less than 0.05 was considered significant.

For statistical comparisons of ACTG1 and TMSB4X mass spectrometry results, to account for potential intra-sample correlations, a mixed-effects model with a random intercept for samples was fitted to the data and used t-tests of contrasts to assess differences between groups (Pinheiro and Bates, 2000). The lmerTest R package (version 3.1-0) was used for the computation (Kuznetsova et al., 2017). A p-value less than 0.05 was considered significant.

### Analysis of single-cell proteomics profiles

For single-cell analysis, we filtered all identifications that were robustly detected at least five cells, leaving a total of 75 proteins (or protein groups). The iBAQ values for proteins in the 30 single cells were normalized to the median of the cells’ mean expression and log_2_-transformed; nondetected values were kept as zeros to avoid imputation artifacts. The empirical Bayes framework available in the *sva* R package (10.18129/B9.bioc.sva) was used to remove the batch covariate, while accounting for the FM1-43 levels as covariate of interest (Johnson et al., 2007; Büttner et al., 2019; Luecken and Theis, 2019). The same framework has been used previously to correct for batch effects in protein mass spectrometry data (Carlyle et al., 2017). The resulting values are referred to as log_2_-normalized iBAQ (niBAQ) units. The relationship between protein expression variance and its average expression was fitted using a log-log cubic smoothing spline with four degrees of freedom; 37 proteins with a higher average expression than a log_2_ niBAQ value of 1.0 and a higher variance than the fit (Figure 4A) were kept for dimensionality reduction. The lower-dimensional manifold (four dimensions) was computed and a trajectory was inferred using the CellTrails R package (Ellwanger et al., 2018) (10.18129/B9.bioc.CellTrails). Protein expression dynamics were calculated by fitting the rolling mean (Haghverdi et al., 2016; Keren-Shaul et al., 2017) with a window size of five of the log2 niBAQ values and the pseudotime axis using a cubic smoothing spline function with 4-degrees of freedom.

### Immunocytochemistry

E15 chicken utricles were dissected in ice-cold Medium 199 containing Hanks’ salts (Thermo Fisher) and were fixed with 4% formaldehyde in PBS at 4°C overnight. Utricles were treated with 0.2 M EDTA in PBS until the otoconia became invisible. For embedding, utricles were equilibrated at 50°C in PBS, followed by incubations in 2.5% and 5% low-melt agarose in PBS for 15 min. Utricles were appropriately positioned while the 5% agarose solution was still hot. Cross-sections from E15 utricles were cut to 80 µm thickness with a vibratome (Leica VT1200). Immunocytochemistry was performed as previously described (Scheibinger et al., 2018) using the following primary antibodies: rabbit anti-TMSB4X (1:250, Proteintech, Rosemont, IL, 19850-1-AP); rabbit anti-AGR3 (1:250, Proteintech, 11967-1-AP); rabbit anti-CRABP1 (1:250, Proteintech, 12588-1-AP); mouse anti-G-actin (1:250, Developmental Studies Hybridoma Bank, JLA20); mouse anti-otoferlin (1:250, Developmental Studies Hybridoma Bank, HCS-1); goat anti-SOX2 (1:100, Santa Cruz Biotechnology, Dallas, TX, sc-17320); mouse anti-tubulin beta-3 (TUJ1. 1:250, BioLegend, San Diego, CA, 801202); rabbit anti-parvalbumin3/ oncomodulin (Heller et al., 2002) (1:1000, Heller laboratory); and rabbit anti-MYO7A (1:1000, Proteus Biosciences, Ramona, CA, 25-6790). Secondary antibodies: Alexa Fluor 546 donkey anti-rabbit (1:250, Thermo Fisher, A10040); Alexa Fluor 647 donkey anti-mouse (1:100, Thermo Fisher, A31571); Alexa Fluor 488 donkey anti-goat (1:250, Thermo Fisher, A11055). Validation of antibodies is reported in Table 2.

**Table 2.** Antibodies used.

| **Antibody** | **Dilution** | **Supplier** | **Catalog #** | **Validation** |
| --- | --- | --- | --- | --- |
| rabbit anti-TMSB4X | 1:250 | Proteintech | 19850-1-AP | (Zhou et al., 2013; Li et al., 2018) |
| rabbit anti-AGR3<br>1:250 | 1:250 | Proteintech | 11967-1-AP | From manufacturer's website: correct-sized band by protein immunoblot in SKOV-3 cells, MCF-7 cells (chicken protein 88% identical to mouse) |
| rabbit anti-CRABP1<br>1:250 | 1:250 | Proteintech | 12588-1-AP | From manufacturer's website: correct-sized band by protein immunoblot in human spleen tissue, transfected HEK-293 cells (chicken protein 95% identical to human) |
| mouse anti-G-actin<br>1:250 | 1:250 | Developmental Studies Hybridoma Bank | JLA20 | (Lin, 1981) |
| mouse anti-otoferlin<br>1:250 | 1:250 | Developmental Studies Hybridoma Bank | HCS-1 | (Goodyear et al., 2010) (chicken protein 73% identical to mouse) |
| goat anti-SOX2<br>1:100 | 1:100 | Santa Cruz Biotechnology | sc-17320 | From manufacturer's website: "recommended for detection of Sox-2 of mouse, rat, human and avian origin by WB, IP, IF, IHC(P) and ELISA" |
| mouse anti-tubulin beta-3 TUJ1. 1:250 | 1:250 | BioLegend | 801202 | (Lee et al., 1990) |
| rabbit anti-parvalbumin3/<br>oncomodulin | 1:1000 | Heller lab | 16910 | (Heller et al., 2002) |
| rabbit anti-MYO7A | 1:1000 | Proteus Biosciences | 25-6790 | (Morgan et al., 2016) |

DAPI (4,6-diamidino-2-phenylindole, 1 µg/ml, Thermo Fisher, D1306) was used to visualize nuclei and Alexa Fluor 488-conjugated phalloidin (1:1000, Invitrogen, Carlsbad, CA, A12379) to visualize F-actin filaments. Sections were imaged with a Plan-Apochromat 40x/1.3 NA oil DIC UV-IRM27 objective on a Zeiss LSM 880 Airyscan laser scanning confocal microscope and Zen Black software. For whole utricle vibratome cross-sections, confocal z-stacks were imaged with the tiling (6% overlap; mode: bounding grid) and stitching function. The 40x Plan-Apochromat objective was used with 0.9x zoom setting. For the extrastriolar and striolar regions, confocal z-stacks were collected separately with the 40x Plan-Apochromat objective used with 1.7 x zoom setting. Extrastriolar and striolar regions in whole utricle vibratome cross-sections were identified using SOX2, TUJ1 and MYO7A antibody labeling as previously described (Ellwanger et al., 2018) (Figure 3—Figure Supplement 6). Maturing or mature striolar type I hair cells express MYO7A but lack SOX2 expression and harbor calyx type terminals. Bouton-innervated extrastriolar Type II hair cells express both, MYO7A and SOX2. All supporting cells express SOX2 and lack MYO7A as well as TUJ1 expression (Figure 3—Figure Supplement 6). Maximum intensity projections were generated by a subset of the z-stacks to preserve single cell resolution.

### Single cell isolation and flow cytometry for RNA-seq

Single cells from E15 chicken utricles were collected and sorted as previously described (Ellwanger et al., 2018). In this study, FM1-43 labeling was not performed and single cells were sorted with a BD FACSAria Fusion instrument (BD Biosciences). Two independent batches of 270 cells were deposited into individual wells of 96-well plates, with prefilled wells of 4 µl lysis solution with 1 U/µl of recombinant RNase inhibitor (Clontech #2313B), 0.1% Triton X-100 (Thermo #85111), 2.5 mM dNTP (Thermo Fisher #10297018), 2.5 µM oligo d(T)30 VN (5’-AAGCAGTGGTATCAACGCAGAGTACT30VN-3’, IDT). Plates containing sorted cells were immediately sealed, frozen on dry ice and stored at −80°C.

### Single-cell RNA-seq

Single-cell RNA-seq was performed via the method of Picelli and colleagues (Picelli et al., 2014) using SMARTscribe (Clontech #639538) for reverse transcription. Kapa Biosystems Hifi HotStart ReadyMix (2X) (#KK2602) was used for 22 cycles of amplification. Amplified cDNAs were purified by bead cleanup using a Biomek FX automated platform and assessed with a fragment analyzer for quantitation and quality assurance. Barcoded libraries were synthesized using a scaled-down Nextera XT protocol (Mora-Castilla et al., 2016) in a total volume of 4 µl. A total of 384 libraries were pooled and paired-end sequenced (2 x 150 bp) on a NextSeq 500/550 High Output flow cell. Raw reads in FASTQ format were aligned to the NCBI Gallus gallus v5.0 (GCA_000002315.3) reference genome using custom scripts on the Sherlock Supercomputer Cluster (Stanford). The FastQC tool (version 0.11.6) was used to run an initial quality control check on the raw sequence data. Sequencing reads were mapped by STAR aligner and the transcriptome BAM files were quantified by RSEM. The results were summarized into counts, fragments per kilobase of transcript per million (FPKM), and transcript per million (TPM) expression matrices.

Scater (10.18129/B9.bioc.scater) was used to perform the quality control of the count expression matrix of 384 cells and 19,153 genes. ERCC spike-in transcripts, 56 information-poor cells and 3,422 low level expressed genes were removed from the count matrix before read count normalization using SCnorm (Bacher et al., 2017). The CellTrails R package (10.18129/B9.bioc.CellTrails) was then utilized following the strategy described in our previous study (Ellwanger et al., 2018). The variable trajFeatureNames was set to the 182 previously used assay genes with the addition of *AGR3*, *AK1*, *GPX2*, and *TMSB4X*, which restricted to the analysis to 186 genes (Table 3).

**Table 3.**
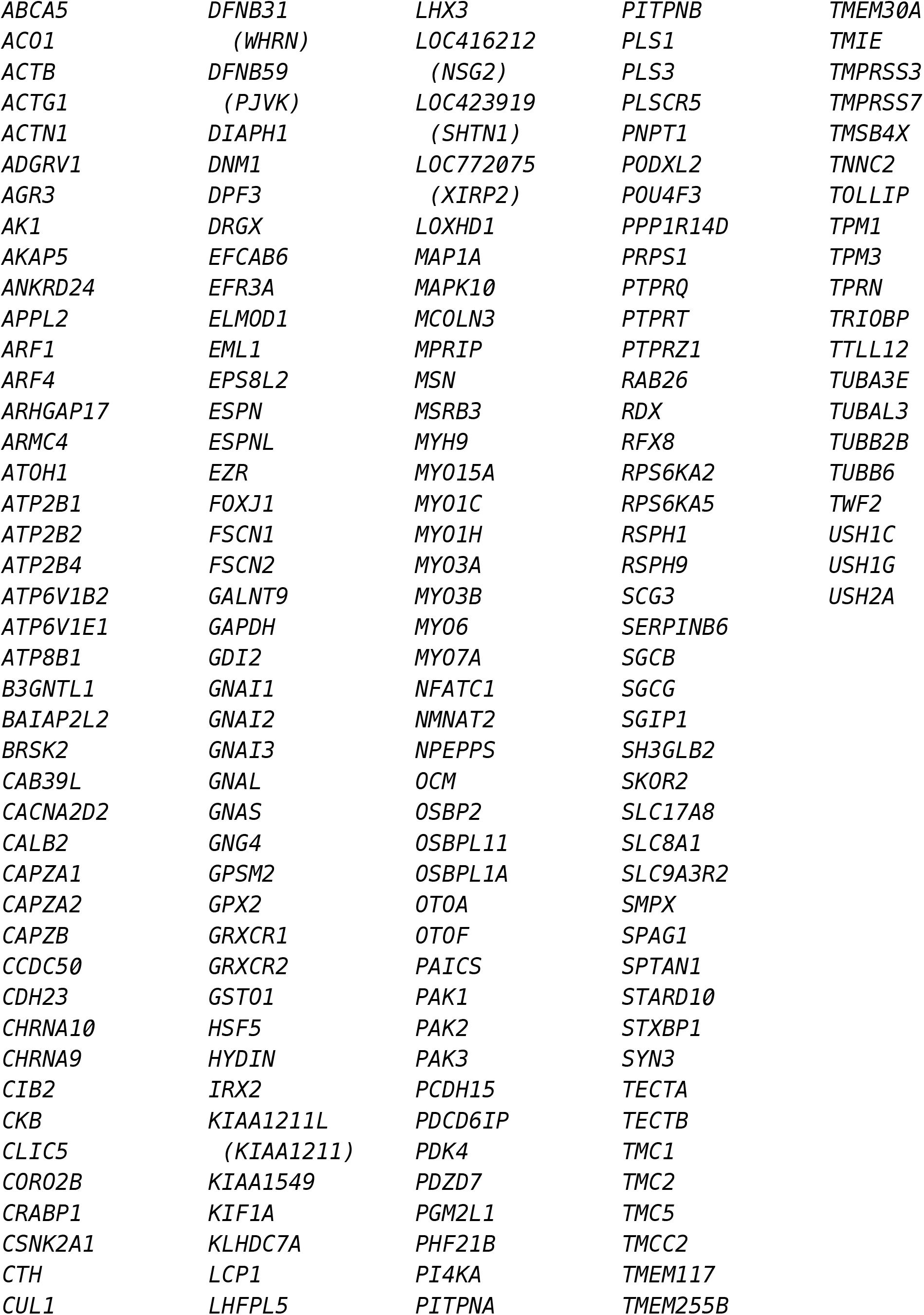
Genes included in scRNA-seq CellTrails analysis.

### Replicates

In all cases, samples were biological replicates—none of the biological samples were split to be run separately as multiple technical replicates. Figure 1. ***B-C***, Confocal imaging of FACS-sorted cells. Experiment was carried out 3 times. ***D-G***, Characterization of mass spectrometry results. Number of samples for FM1-43_high_: 1 cell, *N*=5; 3 cells, *N*=4; 5 cells, *N*=4; 20 cells, *N*=4. Number of samples for FM1-43_low_: 1 cell, *N*=5; 3 cells, *N*=3; 5 cells, *N*=4; 20 cells, *N*=4. ***H-I***, Number of samples for FM1-43_high_: 1 cell, *N*=10; 20 cells, *N*=3. Number of samples for FM1-43_low_: 1 cell, *N*=10; 20 cells, *N*=3. Figure 2. Characterization of FM1-43_high_ and FM1-43_low_ samples. ***A-C***, Experiment was carried out two times. Panels were based on Experiments 1 and 2. Figure 3. ***A-C***, Immunofluorescence detection of AGR3. Experiment was carried out 3 times. ***D-F***, CRABP1. Experiment was carried out 3 times. ***G-I***, TMSB4X. Experiment was carried out 5 times. ***J-L***, G-actin. Experiment was carried out 3 times. ***M-O***, OCM. Experiment was carried out 3 times. Figure 4. Developmental trajectory based on proteomics of single cells. Analysis used 30 single cells from two experiments. Figure 5. Developmental trajectory based on scRNA-seq of single cells. Analysis used two independent batches of 270 cells.

## Data availability

The mass spectrometry proteomics data, including raw data from the mass spectrometry runs, have been deposited to the ProteomeXchange Consortium via the PRIDE partner repository (Perez-Riverol et al., 2019) with the dataset identifier PXD014256. The analyzed data are reported in Figure 1—Source Data 1. The analyzed single-cell RNA-seq data are reported in Figure 5—Source Data 1. The complete analysis of the single-cell RNA-seq will be reported elsewhere.

## Code availability

CellTrails (10.18129/B9.bioc.CellTrails) software is available from Bioconductor (release 3.9). CellTrails is described in detail elsewhere (Ellwanger et al., 2018).

## Figure Supplement Files

**Figure 1—Figure Supplement 1.**
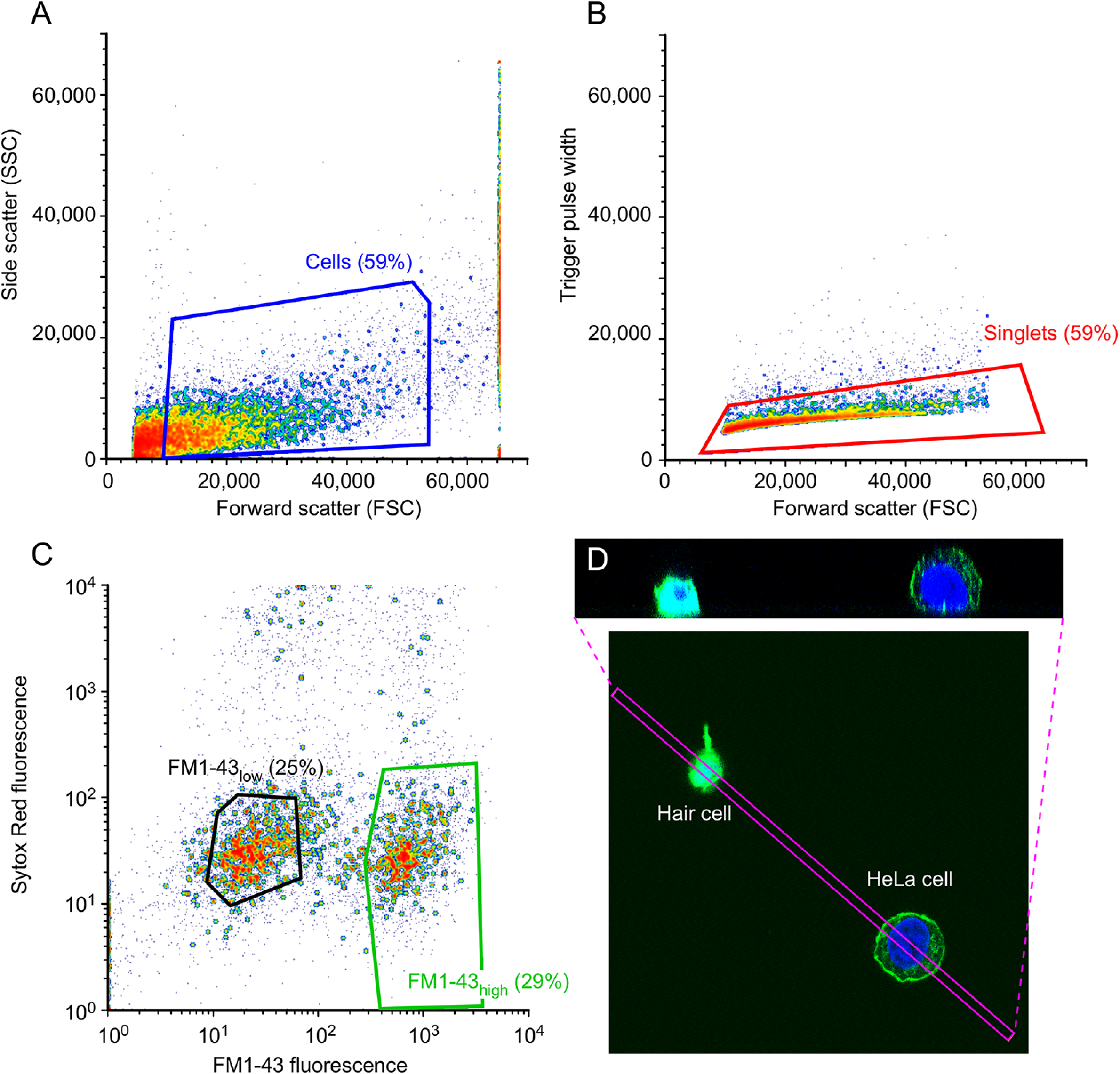
Characterization of isolated utricle cells. ***A-C***, FACS gating protocol. ***D***, Isolated hair cell and HeLa cell, labeled after dissociation with FM1-43 (green) and DAPI (blue). The box indicating the x-y extent of the x-z reslice shown in the upper panel is indicated by the magenta box. Average projection was used.

**Figure 1—Figure Supplement 2.**
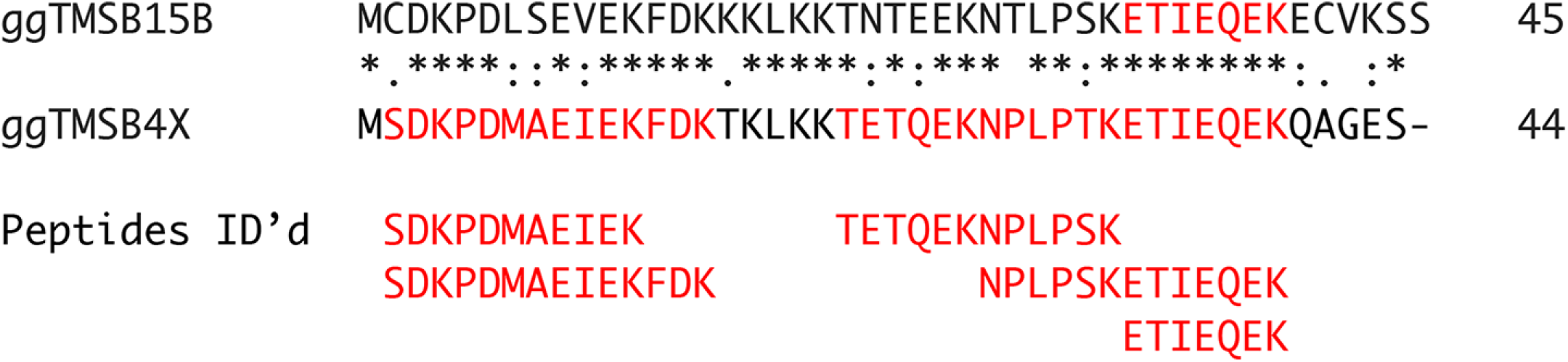
Peptide coverage of TMSB4X in mass spectrometry experiments. Complete sequences for chicken TMSB15B and TMSB4X are aligned, with sequences identified by Andromeda and MaxQuant indicated in red. The actual identified peptides identified are listed below.

**Figure 2—Figure Supplement 1.**
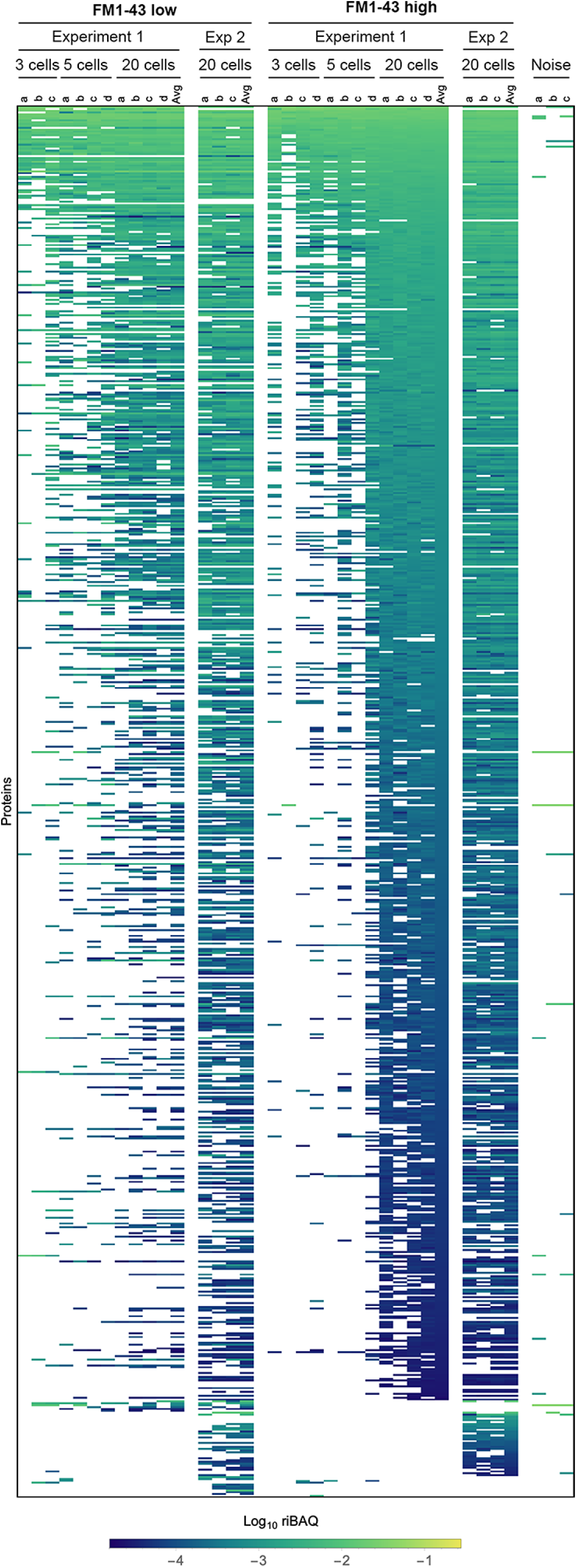
Expression levels for all proteins in samples from Experiments 1 and 2 that contain pools of three cells, five cells, or 20 cells. Each sample (indicated by letters) under the callout for cell numbers contains that number of cells (i.e., on the left, samples a-c under labels indicating “Experiment 1” and “FM1-43 low” each contain three cells). Log10 riBAQ color scale is indicated at bottom. Heat map is sorted by the average of the 20-cell FM1-43_high_ samples from Experiment 1. Three samples (from Experiment 2) that were collected during FACS experiments using noise to gate are also shown.

**Figure 2—Figure Supplement 2.**
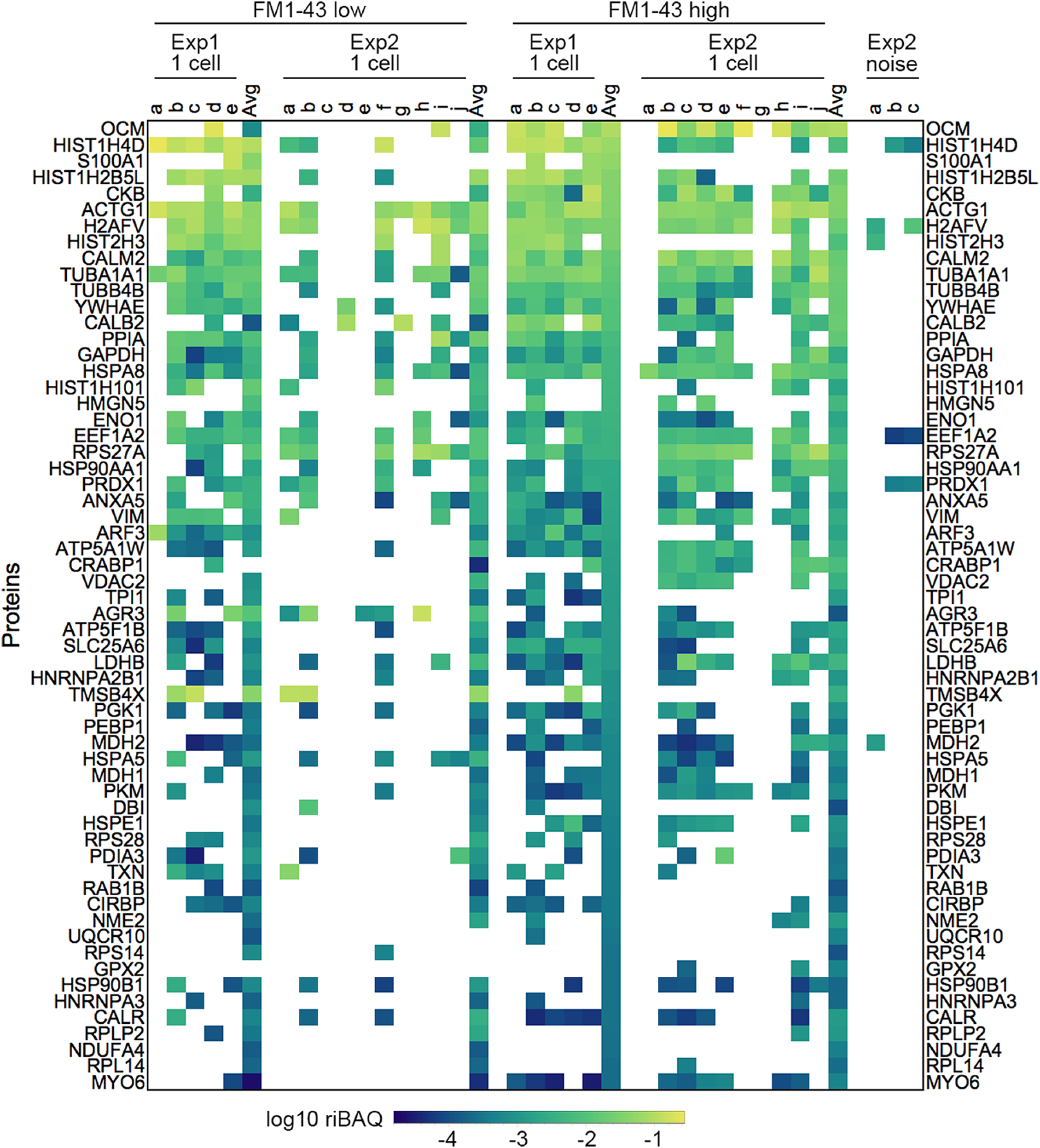
Expression levels for all proteins in samples from Experiments 1 and 2 from single-cell samples. Each sample is indicated by a letters. “Avg” indicates the average of the 20-cell samples for that condition and experiment. Log10 riBAQ color scale is indicated at bottom. Heat map is sorted by the average of the 20-cell FM1-43_high_ samples from Experiment 1. Three samples (from Experiment 2) that were collected during FACS experiments using noise to gate are also shown.

**Figure 2—Figure Supplement 3.**
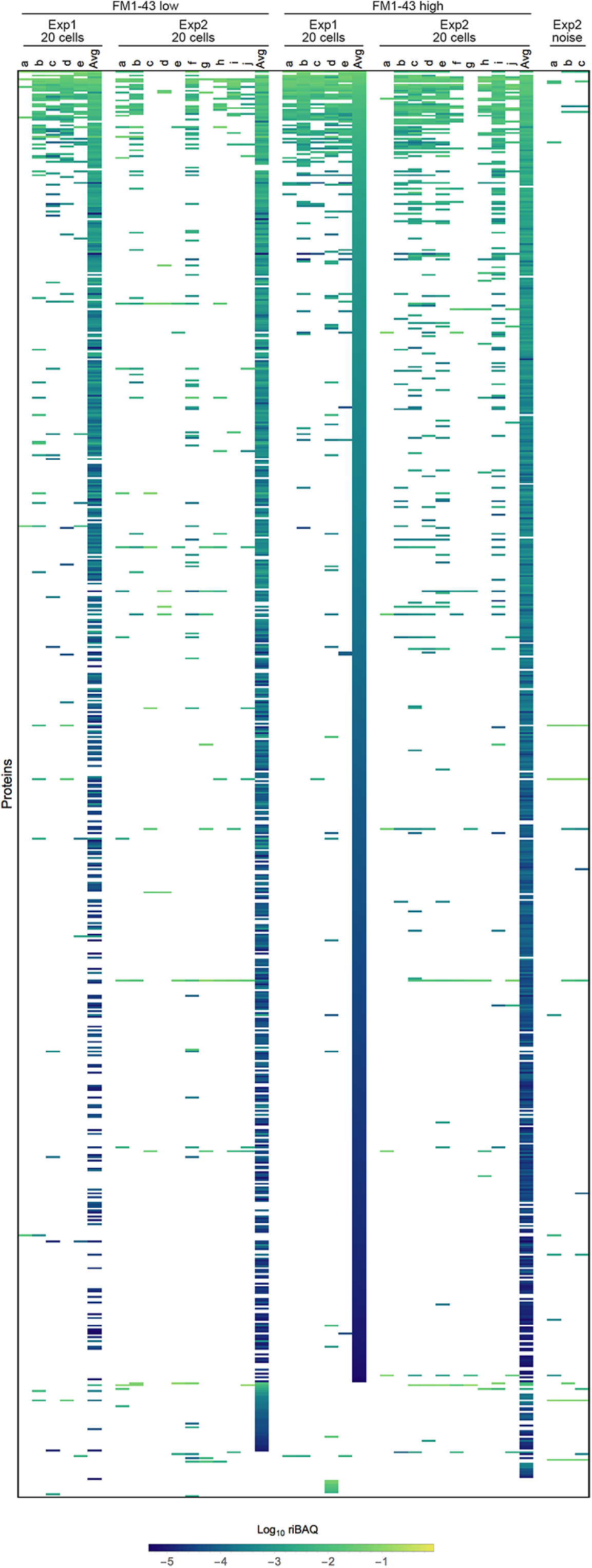
Expression levels for all proteins in single-cell samples in Experiments 1 and 2. Each sample is indicated by a letters. “Avg” indicates the average of the 20-cell samples for that condition and experiment. Heat map, which shows all identified proteins or protein groups in single cells, is sorted by the FM1-43_high_ 20-cell average from Experiment 1. Three samples (from Experiment 2) that were collected during FACS experiments using noise to gate are also shown.

**Figure 3—Figure Supplement 1.**
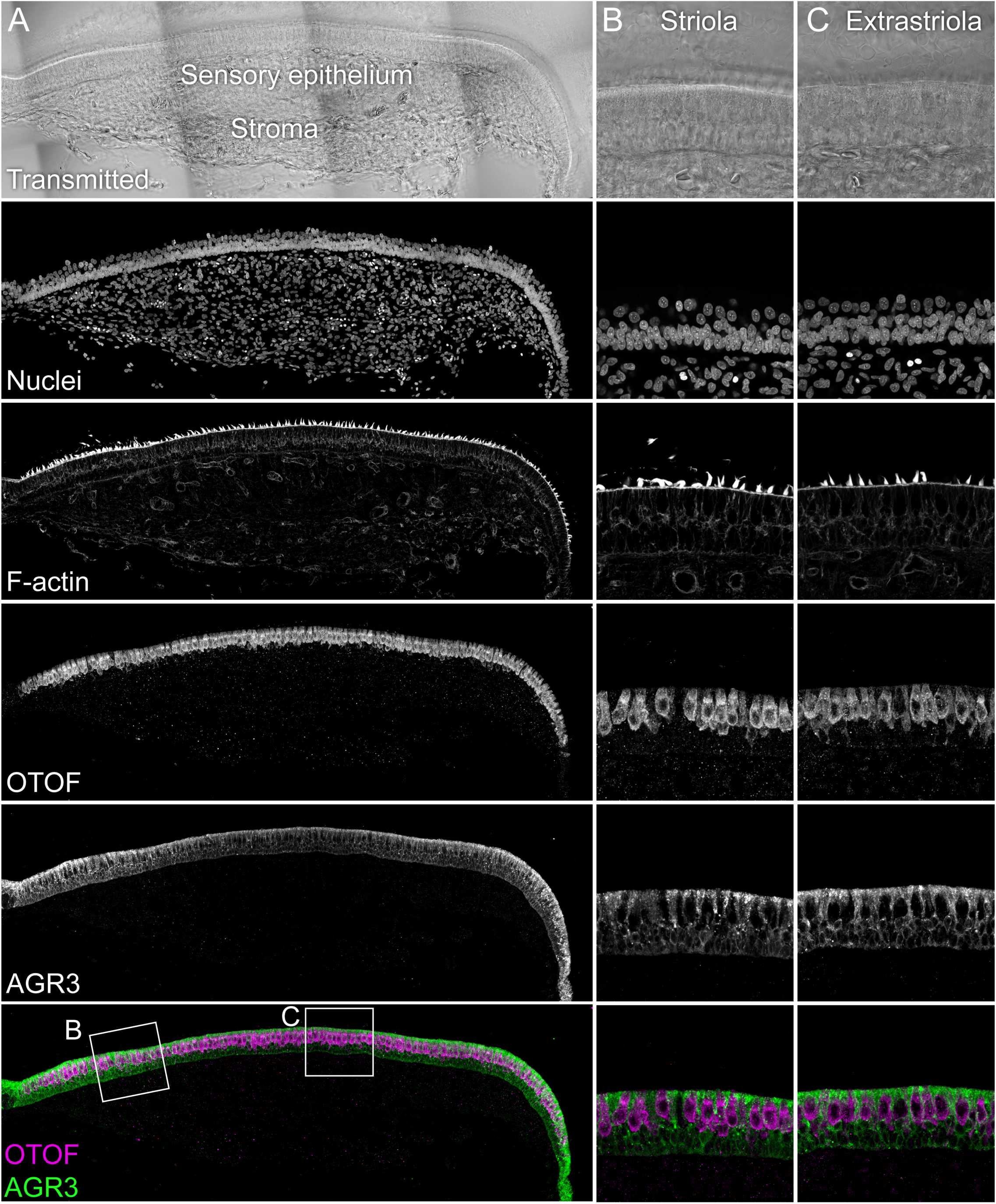
Immunolocalization of AGR3 in E15 chick utricle. The sensory epithelium and the stroma are indicated in the transmitted-light section. Nuclei were stained with DAPI and F-actin with phalloidin; OTOF and AGR3 were stained with specific antibodies. Magnified panels on right clearly show that hair cells (marked by OTOF staining) have much lower levels of AGR3 than do supporting cells, which surround hair cells. Panel full widths: A, 913 µm; B-C, 125 µm.

**Figure 3—Figure Supplement 2.**
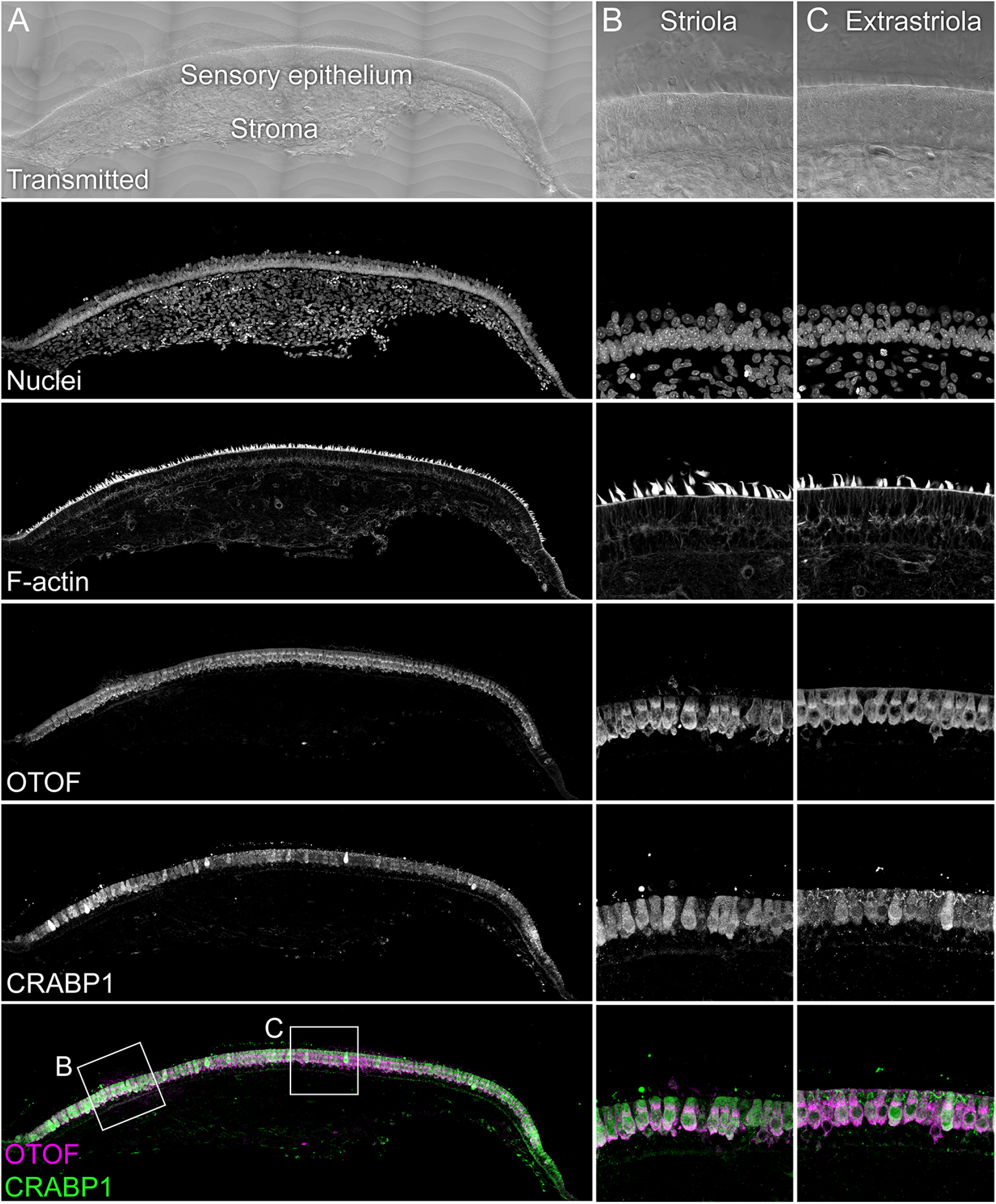
Immunolocalization of CRABP1 in E15 chick utricle. The sensory epithelium and the stroma are indicated in the transmitted-light section. Nuclei were stained with DAPI and F-actin with phalloidin; MYO7A and CRABP1 were stained with specific antibodies. Magnified panels on right clearly show that hair cells are co-labeled by CRABP1 and OTOF antibodies. Panel full widths: A, 1038 µm; B-C, 125 µm.

**Figure 3—Figure Supplement 3.**
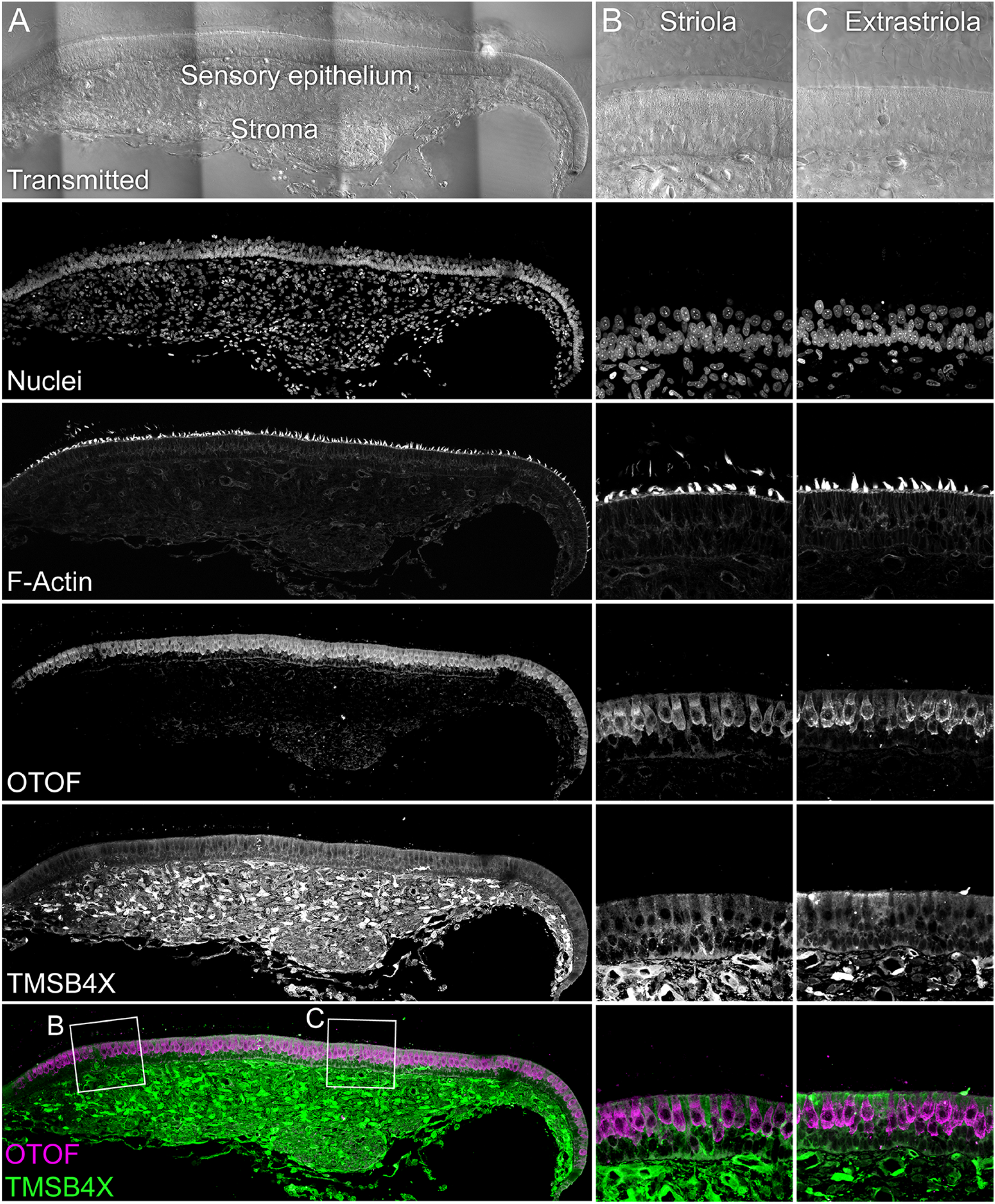
TMSB4X localization. The sensory epithelium and the stroma are indicated in the transmitted-light section. Nuclei were stained with DAPI and F-actin with phalloidin; OTOF and TMSB4X were stained with specific antibodies. TMSB4X is high in the stroma, at moderate levels in supporting cells, and low in hair cells. Panel full widths: A, 934 µm; B-C, 125 µm.

**Figure 3—Figure Supplement 4.**
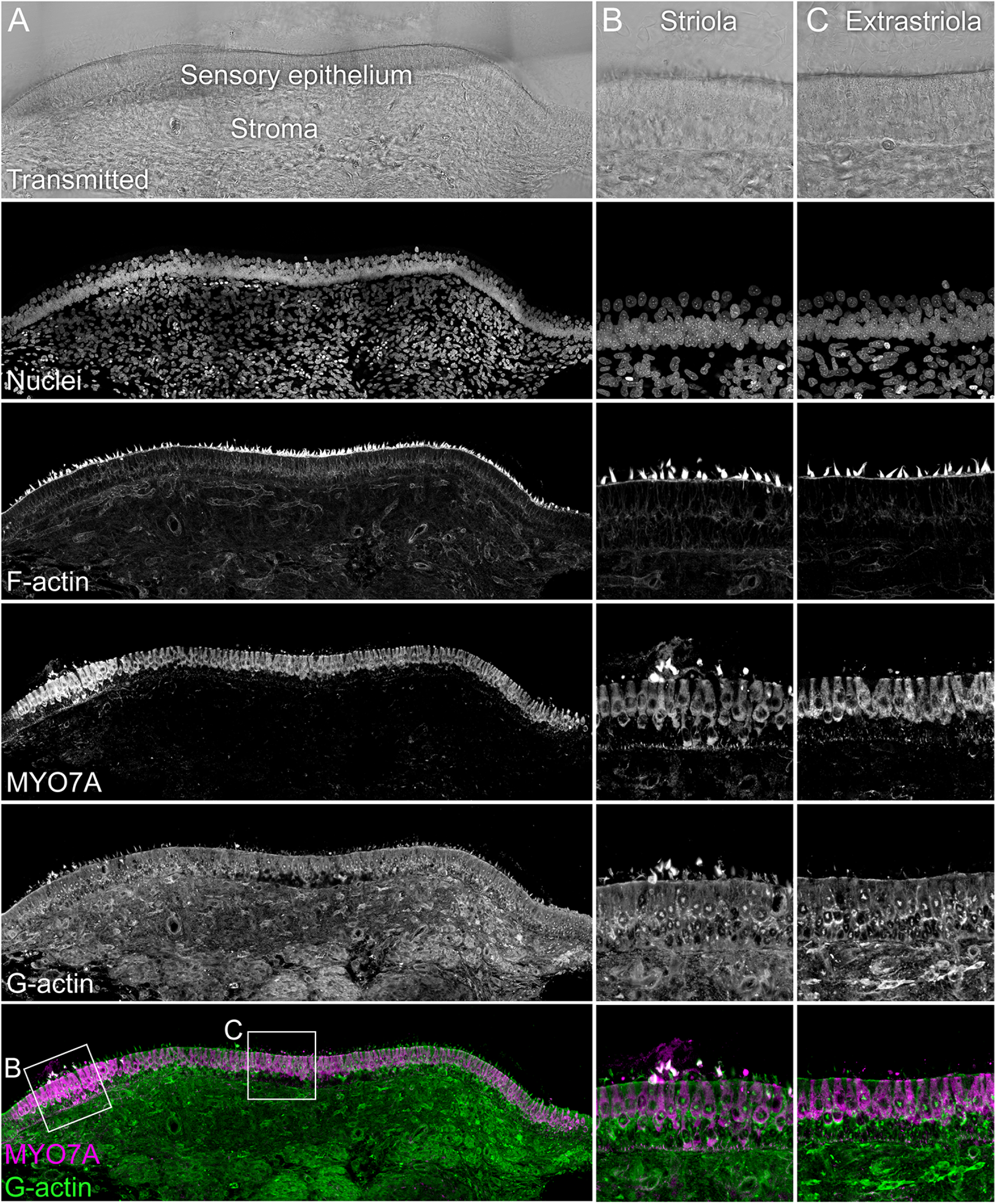
G-actin localization with JLA20 antibody. The sensory epithelium and the stroma are indicated in the transmitted-light section. Nuclei were stained with DAPI and F-actin with phalloidin; MYO7A and G-actin were stained with specific antibodies. Magnified panels on right show that while both hair cells and supporting cells have JLA20 labeling, supporting-cell labeling is more intense. Panel full widths: A, 839 µm; B-C, 125 µm.

**Figure 3—Figure Supplement 5.**
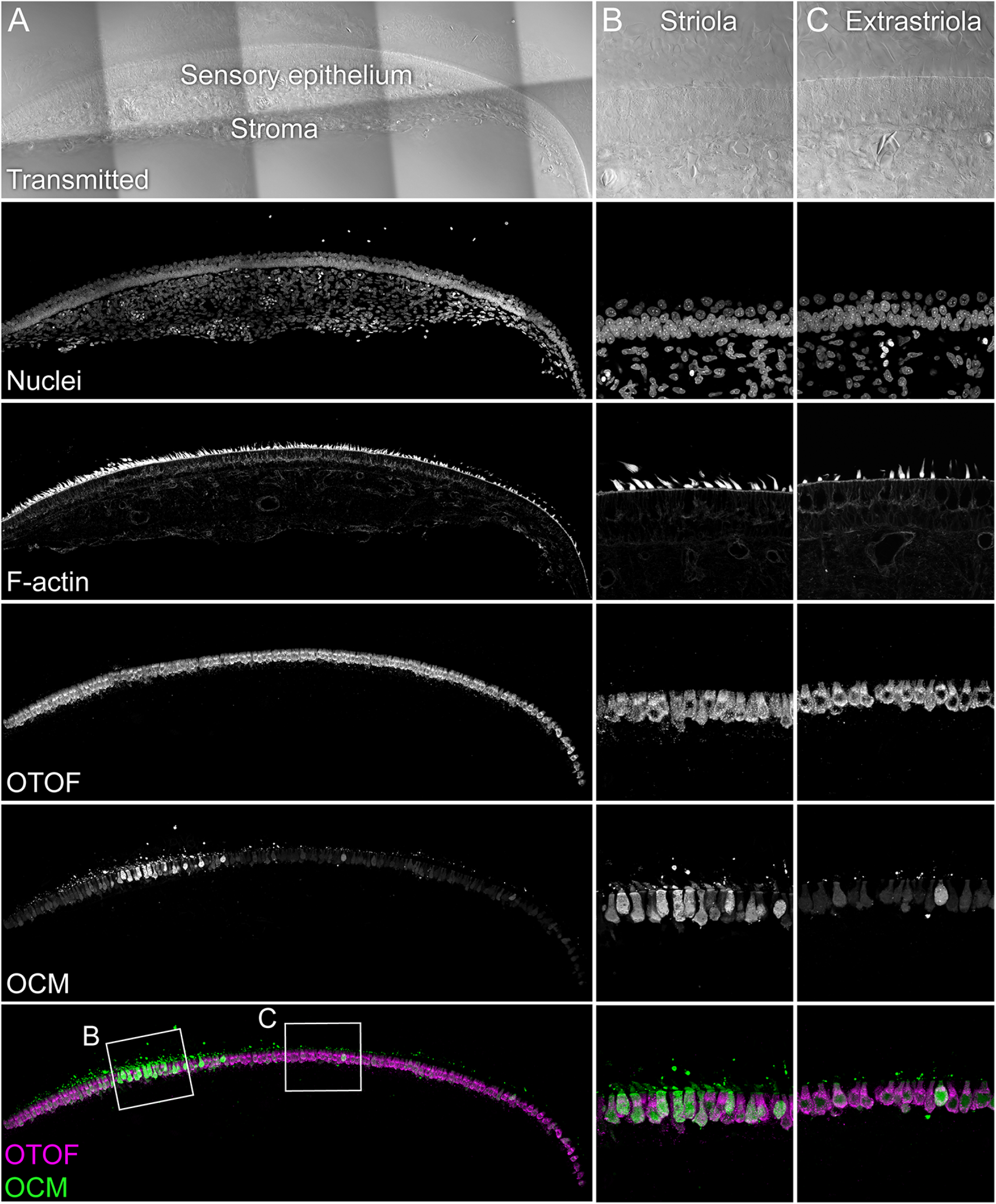
OCM localization with anti-PV3 antibody. The sensory epithelium and the stroma are indicated in the transmitted-light section. Nuclei were stained with DAPI and F-actin with phalloidin; OTOF and OCM were stained with specific antibodies. Magnified panels on right show that OCM labeling is stronger in the striola than the extrastriola, but that occasional extrastriola cells label strongly. Panel full widths: A, 946 µm; B-C, 125 µm.

**Figure 3—Figure Supplement 6.**
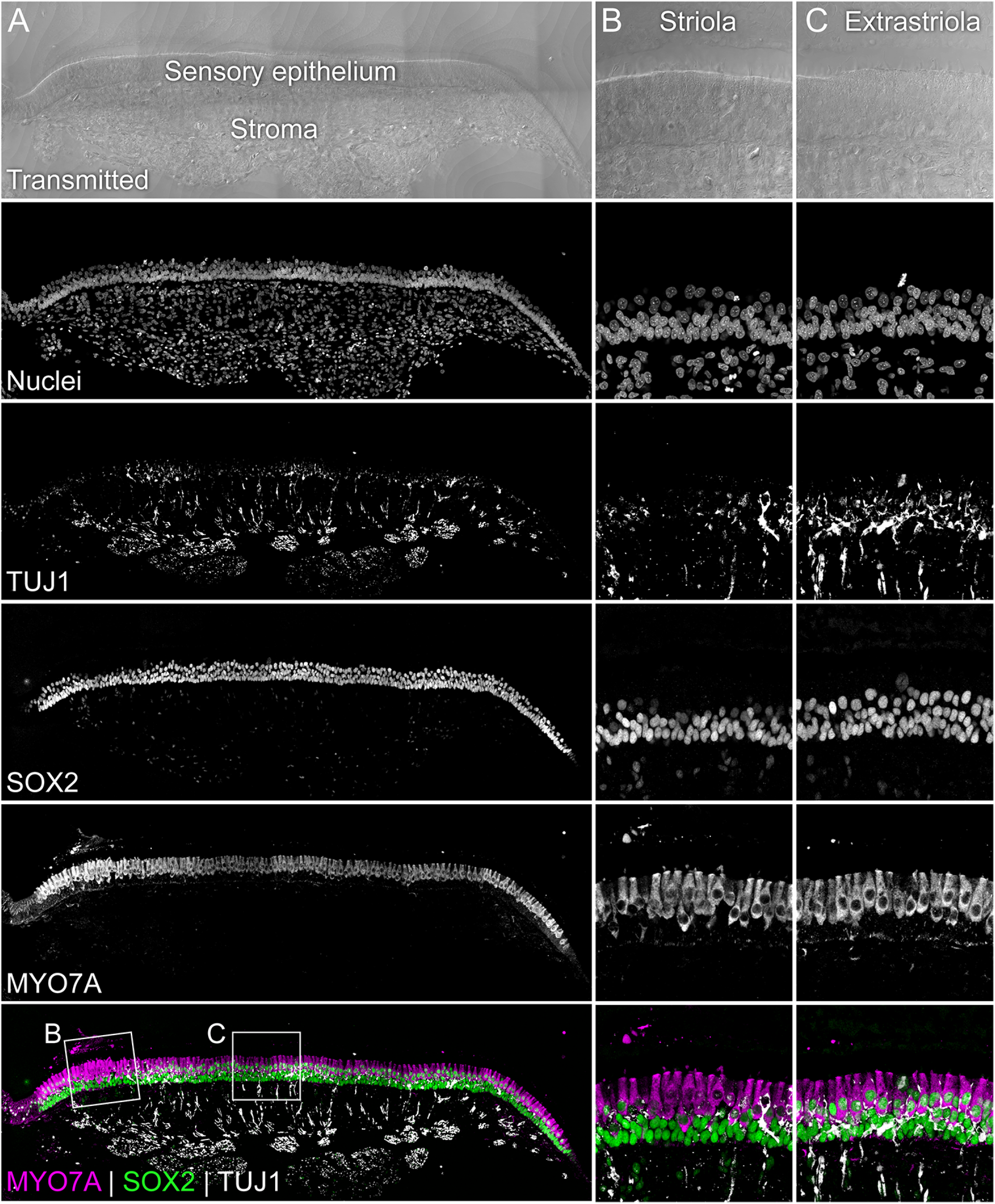
Identification of striola and extrastriola regions. The sensory epithelium and the stroma are indicated in the transmitted-light section. Nuclei were stained with DAPI and F-actin with phalloidin; SOX2, MYO7A and tubulin beta-3 (TUJ1) were stained with specific antibodies. The striolar region harbors maturing/mature type I hair cells (MYO7A^+^/SOX2^-^) with calyx type terminals (TUJ1^+^). Extrastriolar regions and the striola harbor type II hair cells (MYO7A^+^/SOX2^+^), which are innervated by boutons (TUJ1^+^). Panel full widths: A, 982 µm; B-C, 125 µm.

## Source Data Files

Figure 1—Source Data 1. **MaxQuant analysis of single cell proteomics data.** Excel file containing data from Experiments 1 and 2 analyzed together by MaxQuant and Andromeda.

Figure 5—Source Data 1. **CellTrails analysis of single-cell RNA-seq data.** Excel file including: Tab S1A: Single-cell RNA-seq measurements. Count matrix after QC and Normalization (ScNorm) of 186 genes and 328 chicken utricle single cells. Rows display single cells and columns display genes. Non-detected signals were set to 0. Tab S1B: tSNE plots and CellTrails maps. Listed are tSNE plots and CellTrails maps for each gene. The first column lists the gene symbol names, the second column shows the tSNE plots, the third column shows the CellTrails maps with the raw expression values, the fourth column shows the CellTrails maps with smoothed expression values, and the fifth column shows the CellTrails maps with the topographical expression surface. Tab S1C: Inferred expression dynamics. Shown is individual pseudotime gene expression along the extrastriolar (TrES, TrES*) and striolar (TrS) trajectory.

## References

Avenarius, M.R., Saylor, K.W., Lundeberg, M.R., Wilmarth, P.A., Shin, J.B., Spinelli, K.J., Pagana, J.M., Andrade, L., Kachar, B., Choi, D., David, L.L. & Barr-Gillespie, P.G. (2014) Correlation of actin crosslinker and capper expression levels with stereocilia growth phases. Mol Cell Proteomics, 13, 606–620.

Bacher, R., Chu, L.F., Leng, N., Gasch, A.P., Thomson, J.A., Stewart, R.M., Newton, M. & Kendziorski, C. (2017) SCnorm: robust normalization of single-cell RNA-seq data. Nat Methods, 14, 584–586.

Belyantseva, I.A., Perrin, B.J., Sonnemann, K.J., Zhu, M., Stepanyan, R., McGee, J., Frolenkov, G.I., Walsh, E.J., Friderici, K.H., Friedman, T.B. & Ervasti, J.M. (2009) γ-actin is required for cytoskeletal maintenance but not development. Proc Natl Acad Sci U S A, 106, 9703–9708.

Benjamini, Y. & Hochberg, Y. (1995) Controlling the false discovery rate: a practical and powerful approach to multiple testing. J. Roy. Statist. Soc. Ser. B, 57, 289–300.

Brown, G.C. (1991) Total cell protein concentration as an evolutionary constraint on the metabolic control distribution in cells. J Theor Biol, 153, 195–203.

Budnik, B., Levy, E., Harmange, G. & Slavov, N. (2018) SCoPE-MS: mass spectrometry of single mammalian cells quantifies proteome heterogeneity during cell differentiation. Genome Biol, 19, 161.

Büttner, M., Miao, Z., Wolf, F.A., Teichmann, S.A. & Theis, F.J. (2019) A test metric for assessing single-cell RNA-seq batch correction. Nat Methods, 16, 43–49.

Carlier, M.F., Jean, C., Rieger, K.J., Lenfant, M. & Pantaloni, D. (1993) Modulation of the interaction between G-actin and thymosin beta 4 by the ATP/ADP ratio: possible implication in the regulation of actin dynamics. Proc Natl Acad Sci U S A, 90, 5034–5038.

Carlyle, B.C., Kitchen, R.R., Kanyo, J.E., Voss, E.Z., Pletikos, M., Sousa, A.M.M., Lam, T.T., Gerstein, M.B., Sestan, N. & Nairn, A.C. (2017) A multiregional proteomic survey of the postnatal human brain. Nat Neurosci, 20, 1787–1795.

Couvillion, S.P., Zhu, Y., Nagy, G., Adkins, J.N., Ansong, C., Renslow, R.S., Piehowski, P.D., Ibrahim, Y.M., Kelly, R.T. & Metz, T.O. (2019) New mass spectrometry technologies contributing towards comprehensive and high throughput omics analyses of single cells. Analyst, 144, 794–807.

Cox, J. & Mann, M. (2008) MaxQuant enables high peptide identification rates, individualized p.p.b.-range mass accuracies and proteome-wide protein quantification. Nat Biotechnol, 26, 1367–1372.

Cox, J., Neuhauser, N., Michalski, A., Scheltema, R.A., Olsen, J.V. & Mann, M. (2011) Andromeda: a peptide search engine integrated into the MaxQuant environment. J Proteome Res, 10, 1794–1805.

Ellwanger, D.C., Scheibinger, M., Dumont, R.A., Barr-Gillespie, P.G. & Heller, S. (2018) Transcriptional dynamics of hair-bundle morphogenesis revealed with CellTrails. Cell Rep, 23, 2901–2914.e14.

Fulton, A.B. (1982) How crowded is the cytoplasm? Cell, 30, 345–347.

Goldschmidt-Clermont, P.J., Furman, M.I., Wachsstock, D., Safer, D., Nachmias, V.T. & Pollard, T.D. (1992) The control of actin nucleotide exchange by thymosin beta 4 and profilin. A potential regulatory mechanism for actin polymerization in cells. Mol Biol Cell, 3, 1015–1024.

Goodyear, R.J., Gates, R., Lukashkin, A.N. & Richardson, G.P. (1999) Hair-cell numbers continue to increase in the utricular macula of the early posthatch chick. J Neurocytol, 28, 851–861.

Goodyear, R.J., Legan, P.K., Christiansen, J.R., Xia, B., Korchagina, J., Gale, J.E., Warchol, M.E., Corwin, J.T. & Richardson, G.P. (2010) Identification of the hair cell soma-1 antigen, HCS-1, as otoferlin. J Assoc Res Otolaryngol, 11, 573-586.

Haghverdi, L., Büttner, M., Wolf, F.A., Buettner, F. & Theis, F.J. (2016) Diffusion pseudotime robustly reconstructs lineage branching. Nat Methods, 13, 845–848.

Heller, S., Bell, A.M., Denis, C.S., Choe, Y. & Hudspeth, A.J. (2002) Parvalbumin 3 is an abundant Ca2+ buffer in hair cells. J Assoc Res Otolaryngol, 3, 488–498.

Herget, M., Scheibinger, M., Guo, Z., Jan, T.A., Adams, C.M., Cheng, A.G. & Heller, S. (2013) A simple method for purification of vestibular hair cells and non-sensory cells, and application for proteomic analysis. PLoS One, 8, e66026.

Höfer, D., Ness, W. & Drenckhahn, D. (1997) Sorting of actin isoforms in chicken auditory hair cells. J Cell Sci, 110, 765–770.

Johnson, W.E., Li, C. & Rabinovic, A. (2007) Adjusting batch effects in microarray expression data using empirical Bayes methods. Biostatistics, 8, 118–127.

Keren-Shaul, H., Spinrad, A., Weiner, A., Matcovitch-Natan, O., Dvir-Szternfeld, R., Ulland, T.K., David, E., Baruch, K., Lara-Astaiso, D., Toth, B., Itzkovitz, S., Colonna, M., Schwartz, M. & Amit, I. (2017) A Unique Microglia Type Associated with Restricting Development of Alzheimer’s Disease. Cell, 169, 1276–1290.e17.

Krey, J.F., Krystofiak, E.S., Dumont, R.A., Vijayakumar, S., Choi, D., Rivero, F., Kachar, B., Jones, S.M. & Barr-Gillespie, P.G. (2016) Plastin 1 widens stereocilia by transforming actin filament packing from hexagonal to liquid. J Cell Biol, 215, 467–482.

Krey, J.F., Wilmarth, P.A., Shin, J.B., Klimek, J., Sherman, N.E., Jeffery, E.D., Choi, D., David, L.L. & Barr-Gillespie, P.G. (2014) Accurate label-free protein quantitation with high- and low-resolution mass spectrometers. J Proteome Res, 13, 1034–1044.

Kuznetsova, A., Brockhoff PB, & RHB, C. (2017) lmerTest Package: Tests in Linear Mixed Effects Models. Journal of Statistical Software, 82, 1–26.

Lee, M.K., Rebhun, L.I. & Frankfurter, A. (1990) Posttranslational modification of class III beta-tubulin. Proc Natl Acad Sci U S A, 87, 7195–7199.

Li, H., Li, Q., Zhang, X., Zheng, X., Zhang, Q. & Hao, Z. (2018) Thymosin β4 suppresses CCl_4_-induced murine hepatic fibrosis by down-regulating transforming growth factor β receptor-II. J Gene Med, 20, e3043.

Lin, J.J. (1981) Monoclonal antibodies against myofibrillar components of rat skeletal muscle decorate the intermediate filaments of cultured cells. Proc Natl Acad Sci U S A, 78, 2335–2339.

Liu, Y., Beyer, A. & Aebersold, R. (2016) On the Dependency of Cellular Protein Levels on mRNA Abundance. Cell, 165, 535–550.

Luecken, M.D. & Theis, F.J. (2019) Current best practices in single-cell RNA-seq analysis: a tutorial. Mol Syst Biol, 15, e8746.

Mora-Castilla, S., To, C., Vaezeslami, S., Morey, R., Srinivasan, S., Dumdie, J.N., Cook-Andersen, H., Jenkins, J. & Laurent, L.C. (2016) Miniaturization Technologies for Efficient Single-Cell Library Preparation for Next-Generation Sequencing. J Lab Autom, 21, 557–567.

Morgan, C.P., Krey, J.F., Grati, M., Zhao, B., Fallen, S., Kannan-Sundhari, A., Liu, X.Z., Choi, D., Müller, U. & Barr-Gillespie, P.G. (2016) PDZD7-MYO7A complex identified in enriched stereocilia membranes. Elife, 5, e18312. doi: 10.7554/eLife.18312.

Nachmias, V.T. (1993) Small actin-binding proteins: the beta-thymosin family. Curr Opin Cell Biol, 5, 56–62.

Park, K., Jang, J., Irimia, D., Sturgis, J., Lee, J., Robinson, J.P., Toner, M. & Bashir, R. (2008) ‘Living cantilever arrays’ for characterization of mass of single live cells in fluids. Lab Chip, 8, 1034–1041.

Patrinostro, X., Roy, P., Lindsay, A., Chamberlain, C.M., Sundby, L.J., Starker, C.G., Voytas, D.F., Ervasti, J.M. & Perrin, B.J. (2018) Essential nucleotide- and protein-dependent functions of Actb/β-actin. Proc Natl Acad Sci U S A, 115, 7973–7978.

Perez-Riverol, Y., Csordas, A., Bai, J., Bernal-Llinares, M., Hewapathirana, S., Kundu, D.J., Inuganti, A., Griss, J., Mayer, G., Eisenacher, M., Pérez, E., Uszkoreit, J., Pfeuffer, J., Sachsenberg, T., Yilmaz, S., Tiwary, S., Cox, J., Audain, E., Walzer, M., Jarnuczak, A.F., Ternent, T., Brazma, A. & Vizcaíno, J.A. (2019) The PRIDE database and related tools and resources in 2019: improving support for quantification data. Nucleic Acids Res, 47, D442–D450.

Perrin, B.J., Sonnemann, K.J. & Ervasti, J.M. (2010) β-actin and γ-actin are each dispensable for auditory hair cell development but required for stereocilia maintenance. PLoS Genet, 6, e1001158.

Picelli, S., Faridani, O.R., Björklund, A.K., Winberg, G., Sagasser, S. & Sandberg, R. (2014) Full-length RNA-seq from single cells using Smart-seq2. Nat Protoc, 9, 171–181.

Pinheiro, J.C. & Bates, D.M. (2000) Mixed-Effects Models in S and S-PLUS. Springer

Ritchie, M.E., Phipson, B., Wu, D., Hu, Y., Law, C.W., Shi, W. & Smyth, G.K. (2015) limma powers differential expression analyses for RNA-sequencing and microarray studies. Nucleic Acids Res, 43, e47.

Roberson, D.F., Weisleder, P., Bohrer, P.S. & Rubel, E.W. (1992) Ongoing production of sensory cells in the vestibular epithelium of the chick. Hear Res, 57, 166–174.

Scheibinger, M., Ellwanger, D.C., Corrales, C.E., Stone, J.S. & Heller, S. (2018) Aminoglycoside Damage and Hair Cell Regeneration in the Chicken Utricle. J Assoc Res Otolaryngol, 19, 17–29.

Schwanhäusser, B., Busse, D., Li, N., Dittmar, G., Schuchhardt, J., Wolf, J., Chen, W. & Selbach, M. (2011) Global quantification of mammalian gene expression control. Nature, 473, 337–342.

Sekerkova, G., Richter, C.P. & Bartles, J.R. (2011) Roles of the espin actin-bundling proteins in the morphogenesis and stabilization of hair cell stereocilia revealed in CBA/CaJ congenic jerker mice. PLoS Genet, 7, e1002032.

Shin, J.B., Streijger, F., Beynon, A., Peters, T., Gadzala, L., McMillen, D., Bystrom, C., Van der Zee, C.E., Wallimann, T. & Gillespie, P.G. (2007) Hair bundles are specialized for ATP delivery via creatine kinase. Neuron, 53, 371–386.

Shin, J.B., Krey, J.F., Hassan, A., Metlagel, Z., Tauscher, A.N., Pagana, J.M., Sherman, N.E., Jeffery, E.D., Spinelli, K.J., Zhao, H., Wilmarth, P.A., Choi, D., David, L.L., Auer, M. & Barr-Gillespie, P.G. (2013) Molecular architecture of the chick vestibular hair bundle. Nat Neurosci, 16, 365–374.

Shin, J.B., Longo-Guess, C.M., Gagnon, L.H., Saylor, K.W., Dumont, R.A., Spinelli, K.J., Pagana, J.M., Wilmarth, P.A., David, L.L., Gillespie, P.G. & Johnson, K.R. (2010) The R109H variant of fascin-2, a developmentally regulated actin crosslinker in hair-cell stereocilia, underlies early-onset hearing loss of DBA/2J mice. J Neurosci, 30, 9683–9694.

Srivastava, D.K. & Bernhard, S.A. (1986) Enzyme-enzyme interactions and the regulation of metabolic reaction pathways. Curr Top Cell Regul, 28, 1–68.

Sun, H.Q., Kwiatkowska, K. & Yin, H.L. (1995) Actin monomer binding proteins. Curr Opin Cell Biol, 7, 102–110.

Tilney, L.G. & Tilney, M.S. (1988) The actin filament content of hair cells of the bird cochlea is nearly constant even though the length, width//number of stereocilia vary depending on the hair cell location. J. Cell Biol., 107, 2563–2574.

Tilney, L.G., Tilney, M.S. & DeRosier, D.J. (1992) Actin filaments, stereocilia, and hair cells: how cells count and measure. Ann. Rev. Cell Biol., 8, 257–274.

Tyanova, S., Temu, T. & Cox, J. (2016) The MaxQuant computational platform for mass spectrometry-based shotgun proteomics. Nat Protoc, 11, 2301–2319.

Weber, A., Nachmias, V.T., Pennise, C.R., Pring, M. & Safer, D. (1992) Interaction of thymosin beta 4 with muscle and platelet actin: implications for actin sequestration in resting platelets. Biochemistry, 31, 6179–6185.

Weydert, S., Zürcher, S., Tanner, S., Zhang, N., Ritter, R., Peter, T., Aebersold, M.J., Thompson-Steckel, G., Forró, C., Rottmar, M., Stauffer, F., Valassina, I.A., Morgese, G., Benetti, E.M., Tosatti, S. & Vörös, J. (2017) Easy to Apply Polyoxazoline-Based Coating for Precise and Long-Term Control of Neural Patterns. Langmuir, 33, 8594–8605.

Wilmarth, P.A., Krey, J.F., Shin, J.B., Choi, D., David, L.L. & Barr-Gillespie, P.G. (2015) Hair-bundle proteomes of avian and mammalian inner-ear utricles. Sci Data, 2, 150074.

Zhao, L., Kroenke, C.D., Song, J., Piwnica-Worms, D., Ackerman, J.J. & Neil, J.J. (2008) Intracellular water-specific MR of microbead-adherent cells: the HeLa cell intracellular water exchange lifetime. NMR Biomed, 21, 159–164.

Zhou, Y., Li, S., Huang, Q., Xie, L. & Zhu, X. (2013) Nanog suppresses cell migration by downregulating thymosin β4 and Rnd3. J Mol Cell Biol, 5, 239–249.

Zhu, Y., Clair, G., Chrisler, W.B., Shen, Y., Zhao, R., Shukla, A.K., Moore, R.J., Misra, R.S., Pryhuber, G.S., Smith, R.D., Ansong, C. & Kelly, R.T. (2018a) Proteomic Analysis of Single Mammalian Cells Enabled by Microfluidic Nanodroplet Sample Preparation and Ultrasensitive NanoLC-MS. Angew Chem Int Ed Engl, 57, 12370–12374.

Zhu, Y., Piehowski, P.D., Zhao, R., Chen, J., Shen, Y., Moore, R.J., Shukla, A.K., Petyuk, V.A., Campbell-Thompson, M., Mathews, C.E., Smith, R.D., Qian, W.J. & Kelly, R.T. (2018b) Nanodroplet processing platform for deep and quantitative proteome profiling of 10-100 mammalian cells. Nat Commun, 9, 882.

Zhu, Y., Podolak, J., Zhao, R., Shukla, A.K., Moore, R.J., Thomas, G.V. & Kelly, R.T. (2018c) Proteome Profiling of 1 to 5 Spiked Circulating Tumor Cells Isolated from Whole Blood Using Immunodensity Enrichment, Laser Capture Microdissection, Nanodroplet Sample Processing, and Ultrasensitive nanoLC-MS. Anal Chem, 90, 11756–11759.

Zhu, Y., Zhao, R., Piehowski, P.D., Moore, R.J., Lim, S., Orphan, V.J., Paša-Tolić, L., Qian, W.J., Smith, R.D. & Kelly, R.T. (2018d) Subnanogram proteomics: impact of LC column selection, MS instrumentation and data analysis strategy on proteome coverage for trace samples. Int J Mass Spectrom, 427, 4–10.

